# Discovery of a small molecule that selectively destabilizes Cryptochrome 1 and enhances life span in p53 knockout mice

**DOI:** 10.1101/2021.07.21.453206

**Authors:** Seref Gul, Yasemin Kubra Akyel, Zeynep Melis Gul, Safak Isin, Tuba Korkmaz, Saba Selvi, Ibrahim Danis, Ozgecan Savlug Ipek, Fatih Aygenli, Ali Cihan Taskin, Nuri Ozturk, Narin Ozturk, Durişehvar Özer Ünal, Mustafa Guzel, Metin Turkay, Alper Okyar, Ibrahim Halil Kavakli

## Abstract

Cryptochromes are negative transcriptional regulators of the circadian clock in mammals. It is not clear how reducing the level of endogenous level of the CRY1 in mammals will affect circadian rhythm and the relation of such a decrease with apoptosis is unknown. Here, we discovered a molecule that destabilizes Cryptochrome 1 (CRY1) both *in vitro* and *in vivo*. The small molecule, called M47, selectively enhanced the degradation rate of CRY1 by increasing its ubiquitination and the period of U2OS *Bmal1*-d*Luc* cells. In addition, subcellular fractionation studies from mice liver indicated that M47 enhanced degradation rate of the CRY1 level in the nucleus. Furthermore, M47-mediated CRY1 reduction enhanced cisplatin-induced apoptosis in Ras-transformed p53 null fibroblast cells. Finally, systemic repetitive administration of M47 increased the median lifespan of p53^−/−^ mice by ~25%. Collectively our data suggest that M47 is a very promising molecule to treat forms of cancer depending on the p53 mutation.

## Introduction

The circadian clock generates a 24-hour rhythm through which physiology and behavior adapt to daily changes in the environment ^1^. Many biological processes like hormone secretion and sleep-wake cycles are controlled by the circadian clock ^2^. Therefore, an innate malfunctioning of the circadian clock and the related pathways can cause various pathologies such as sleep disorders, altered metabolism, obesity, diabetes, mood disorders, cancer, and cardiovascular diseases ^3–5^.

At the molecular level, the clockwork of the cell involves several proteins that participate in positive and negative transcriptional/translational feedback loops (TTFL). BMAL1 and CLOCK are transcription factors that bind E-box elements (CACGTG) in clock-controlled genes including *Period* and *Cryptochrome* and thereby exert a positive effect on the circadian transcription ^6, 7^. The mammalian PERIOD (PER) and CRYPTOCHROME (CRY) proteins form heterodimers that interact with casein kinase Iε (CKIε) and then translocate into the nucleus. Here, where CRY acts as a negative regulator of BMAL1/CLOCK–driven transcription ^8^. Upon phosphorylation, CRYs are ubiquitinated by E3 ubiquitin ligases e.g. FBXL3 and FBXL21 are directed to the proteasome for degradation. FBXL3 and FBXL21 act antagonistically on CRY to regulate its stability differentially in the cytosol and nucleus ^9, 10^.

A second feedback loop consists of retinoic acid receptor-related orphan receptors (RORs) and REV-ERBs which control the transcription of the *Bmal1* gene and, in turn, regulate the molecular clock ^7^. The recent studies indicated that 50% of detected metabolites are under circadian control in the mouse liver and nearly 50% of transcripts in at least one tissue are under the control of the circadian clock ^11, 12^. Although circadian clock disruption is related to various diseases, loss of *Cry1* and *Cry2* caused an unexpected effect on cancer and did not aggravate the radiation-induced tumor growth and mortality ^13^. Furthermore, in the background of cancer-prone (*p53*^−/−^) mice, the absence of *Cry*s (*Cry1^−/−^ Cry2^−/−^*) extended the median lifespan 1.5 fold compared to control littermates ^14^. These findings evoked the idea that small molecules destabilizing the CRYs can be used as anti-cancer agents in some cancer types.

Several studies have been carried out to find small molecules to treat diseases associated with the circadian rhythm. These studies used a high-throughput screening assay to look at the phenotypic changes in the circadian rhythm of reporter cells. Multiple molecules that affect various features of circadian rhythm have been identified ^15–18^. Another small molecule, KL001, increased the stability of CRYs and suppressed gluconeogenesis ^17^. Isoform selective KL101 and TH301 molecules stabilizing CRY1 and CRY2, respectively, were shown to regulate the differentiation of brown adipose tissue ^19^. ROR nuclear receptor agonist, nobiletin, improved the amplitude in mice with metabolic syndrome ^20^. Alternatively, a structure-based drug design approach can be utilized to design small molecules by using the available crystal structures of core clock proteins ^21–23^. For example, a molecule (named CLK8) identified from virtual screening was shown to enhance the amplitude of the circadian rhythm by reducing the nuclear levels of CLOCK protein ^16^. In this study, we aimed to find a CRY-binding small molecule using a structure-based approach by targeting the FAD-binding pocket which is critical for the FBXL3 interaction.

We started by virtually screening a library of approximately 2 million small molecules and gradually narrowed down the number of selected molecules by employing different biochemical and molecular evaluation steps. Finally, we identified a CRY-binding small molecule (M47) that destabilizes CRY1 and increases the period length of the circadian rhythm at the cellular level. Further in vivo studies showed that M47 has a half-life of 6h in blood and destabilizes CRY1 in the liver cells of mice. We further showed M47 enhances apoptosis in a Ras-transformed p53-null mouse skin fibroblast cell line. Finally, M47 was administered to p53^−/−^ mice to determine its effect on the lifespan of cancer-prone mice. Our results indicated that p53^−/−^ mice treated with M47 had ~25% increased lifespan compared to control animals. The mild toxicity profile, pharmacokinetic and pharmacodynamics properties of the M47 from mice and in vitro studies suggested that it may be used to improve the effectiveness of cancer treatment-related to p53 mutations and other types of diseases related to CRY1.

## Results

### Structure-Based Small Molecule Design

The mouse CRY1 (mCRY1) structure (PDB ID: 4K0R) was used to identify small molecules targeting its photolyase (PHR) domain. To obtain the receptor (CRY1) at the physiological conditions, the molecular dynamics (MD) simulation was performed on the homology model of the mCRY1 structure based on the 4K0R. Firstly, the structure of mCRY1 was solvated in a rectangular box and neutralized with counterions. Subsequently, the system was minimized, heated up to physiological temperature (310°K), and simulated for 50 ns. The root-mean-square deviation (RMSD) of backbone atoms showed convergence after initial increase upon minimization (Fig. S1). Equilibrated CRY1 structure has been used as the receptor for the docking simulations. A freely available library from Ambinter containing 8 million small molecules with the non-identified function was used as ligands. The library was initially filtered according to “Lipinski’s Rule of Five” ^24^ and the remaining ~1 million small molecules were screened against the PHR domain of CRY1 using AutoDock Vina. PHR domain is critical for the period regulation through CRY1 and interacts with FBXL3, an important ubiquitin ligase for the degradation of CRY1 ^9, 10, 23^. The top 200 compounds based on their binding energies (range from −9 to −13.5 kcal/mol) to the FAD region were selected for further experimental studies (data not shown).

### Identification of the Molecules Effecting the Half-Life of the CRY

Initially, the toxicity of selected molecules was assessed by MTT assay using immortalized human osteosarcoma (U2OS) cells at concentrations of 20, 10 and, 2.5 µM. Molecules with relative cell viability less than 90% at 2.5 µM were excluded from further studies. Molecules changing the stability of CRYs were shown to affect the period length of circadian rhythm ^17, 25^. We, therefore, treated U2OS cells, stably expressing *Bmal1-*d*luc* (where the destabilized form of *Luciferase* gene fused with *Bmal1* promotor), with the nontoxic small molecules and monitored bioluminescence to identify molecules changing the period. We identified a set of molecules affecting the circadian rhythm of U2OS cells either by lengthening or shortening the period by1 hour or more. The effects of these molecules on the half-life of CRY1 were determined using we used CRY1::dLUC degradation assay ^17^ where CRY1 was fused with destabilized Luciferase (dLUC) at the C-terminal. Among CRY1 modulating molecules, M47 showed the best dose-responsive effect (data not shown) and was selected for further characterization.

### Characterization of M47

Since we are mostly using luciferase-based reporter assays, initially we tested the effect of M47 (Fig. 1A) on the half-life of dLUC and then its toxicity on U2OS cells in a dose-dependent manner. M47 did not affect the stability of LUC and showed any toxic effect on the U2OS cell line (Fig. S2A, B). We then determined the effect of M47 on the period of the circadian rhythm using U2OS *Bmal1*-d*Luc* cells and NIH 3T3 *Bmal1-dLuc* cells. M47 significantly lengthened the period in a dose-dependent manner (Fig. 1B). M47 treated, which has no toxic effect on NIH 3T3 cell line, cells exhibited similar phenotype as well (Fig. S2C and D). To eliminate the possibility that M47 might affect the other proteins and, in turn, circadian rhythmicity we tested the effect of M47 in *Cry1^−/−^Cry2^−/−^* mouse embryonic fibroblast cells (DKO-MEF), which has different genetic background, transfected with *mPer2-luc* plasmid as described in ^26^. Results indicated that M47 didn’t affect on luminance dose dependently in the absence of the CRYs, suggesting that the effect of M47 was through CRY (Fig. 1C). We next determined the effect of the M47 on the half-life of CRY1 by measuring the decay rate of CRY1::dLUC and CRY2::dLUC as described in ^17^. To this end, HEK 293T cells were transfected with *Cry-Luc* plasmids, treated with M47 doses, then, treated with cycloheximide to inhibit protein translation. Notably, M47 reduces the half-life of CRY1 in a dose-dependent manner while there was no effect on the half-life of CRY2 (Fig. 2A, B). To verify the CRY1 dependency of M47, we generated U2OS *CRY1^−/−^ Bmal1*-d*Luc* cells by utilizing CRISPR/Cas9 technology (Fig. S2E). Knocking out the *CRY1* in this cell line resulted in a shorter period compared to wild-type controls as in agreement with previously published data ^27^. Notably, when U2OS *CRY1^−/−^ Bmal1*-d*Luc* cells were treated with the M47, no change was observed in the period length of the circadian rhythm (Fig. 2C). Collectively, CRY degradation and circadian phenotypes of *CRY1^−/−^* and wild type U2OS cells showed that the M47 selectively binds to the CRY1 and exerts its effect through CRY1.

**Fig. 1.**
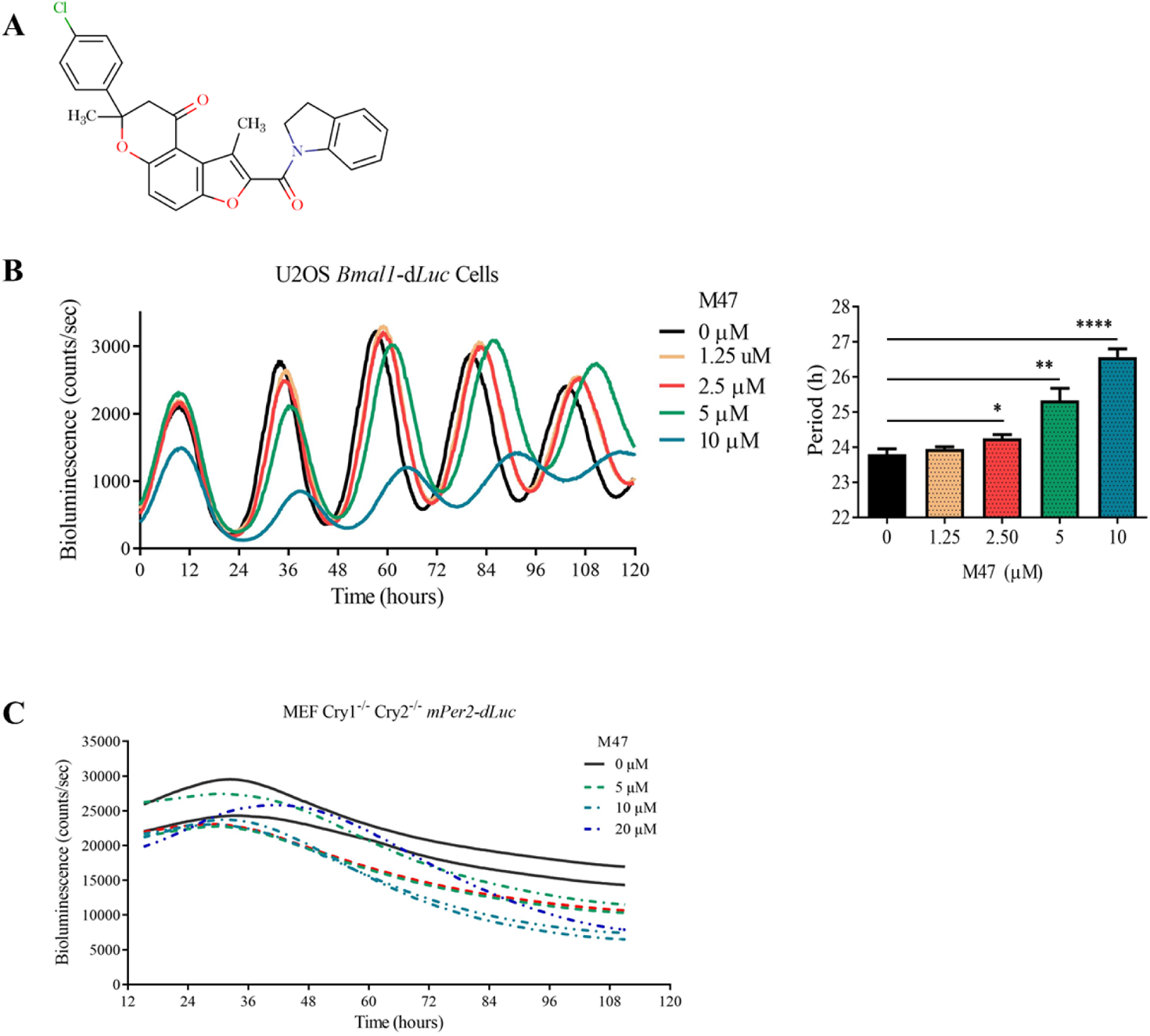
M47 changes period of U2OS cells dose dependently. (**A**) Structure of M47. (**B**) Representative figure for the effect of M47 on the bioluminescence rhythm of U2OS cells stably expressing *Bmal1*-d*Luc*. M47 lengthened the circadian rhythm dose dependently (Data represent the mean ± SEM, n=4 with duplicates ***: p < 0.001 **: p < 0.005, *: p <0.05 versus DMSO control by student t-test). (**C**) Effect of M47 on the *Cry1^−/−^/Cry2^−/−^*mouse embryonic fibroblasts transiently transfected with *mPer2-dLuc* reporter (n=2 with duplicates).

**Fig. 2.**
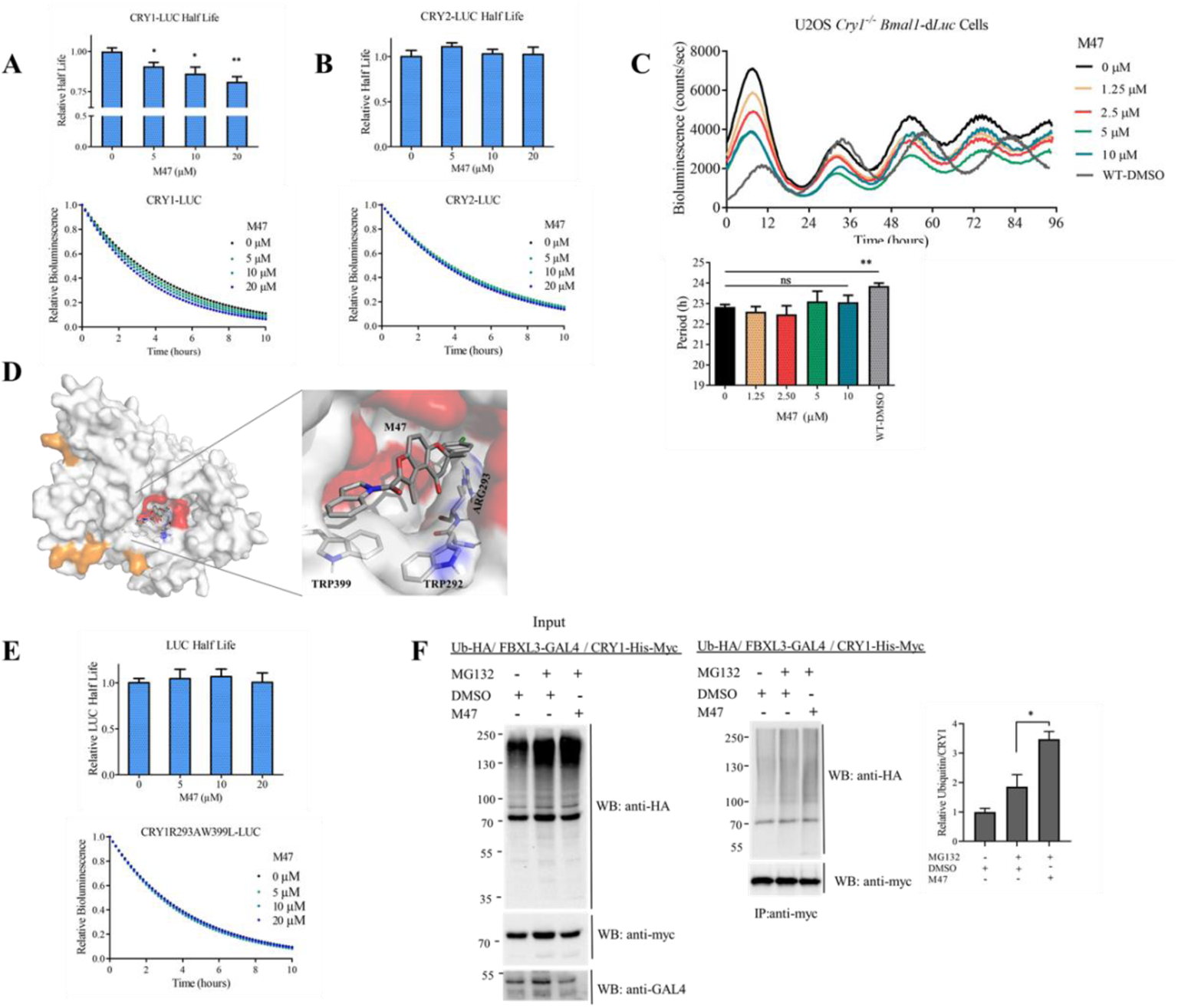
M47 decreases the half-life of CRY1. (**A**) M47 increased the degradation rate of CRY1-LUC dose dependently. HEK 293T cells were transfected with *Cry1-Luc* plasmid. 24h after transfection cells were treated with different doses of M47 or solvent (DMSO final 0.5%) as control. 24h after molecule treatment cells were treated with cycloheximide (20µg/ml final) and bioluminescence was recorded. Normalized half-life is shown with ± SEM (n=4 with triplicates). *p<0.05,**p<0.01 (**B**) M47 did not affect the degradation rate of CRY2-LUC (n=4 with triplicates). (**C**) Representative figure for the effect of M47 on the bioluminescence rhythm of U2OS *CRY1^−/−^ Bmal1*-d*Luc*. U2OS *CRY1^−/−^ Bmal1*-d*Luc* had shorter period length compared to U2OS *Bmal1*-d*Luc.* M47 had no effect on the circadian rhythm of the U2OS *CRY1^−/−^ Bmal1*-d*Luc* in dose dependently (Data represent the mean ± SEM, n=3 with duplicates **: p < 0.005, versus DMSO control by student t-test; ns0statistically not significant). (**D**) Binding pose of M47 on the equilibrated CRY1 (4K0R) structure predicted by AutodockVina with −13.4kcal/mol binding energy. Protein structure is shown in surface. FBXL3 binding residues are colored as red, PER2 binding residues are colored as orange. M47 interacting residues are shown in sticks (carbons are white, nitrogens are blue, oxygens are red, however, carbons in M47 are shown in gray). (**E**) M47 did not affect the half-life of mutant CRY1R293AW399L-LUC degradation (Data shown with ± SEM n=3 with triplicates). **(F)** The ubiquitination of the CRY1 in the presences of M47. The results are the average of 3 biological replicates. (student’s t-test *p<0.05).

The docking pose of M47 showed that it binds to the FAD-binding region of the CRY1 by interacting with R293 and W399 amino acid residues (Fig. 2D). Indoline groups of M47 and Trp399 interact through a strong pi-pi interaction and the benzene group at the other side of the M47 interacts with Arg293 through a pi-cation interaction (Fig. 2D). We hypothesized that the replacement of these residues in CRY1 should eliminate the binding of M47 to CRY1. To test this, R293 and W399 of CRY1 were replaced with Ala and Leu by site-directed mutagenesis, respectively. Before measuring the effect of M47 on this CRY1 mutant, we confirmed CRY1-R293A-W399L mutant retained its repressor activity with *Per1-*d*Luc* assay using BMAL1/CLOCK in the presenceof the mutant and wild type CRY1 (Fig. S3A). We then measured the half-life of CRY1-R293A-W399L::LUC in the presence of M47. M47 did not affect the half-life of CRY1-R293A-W399L compared to DMSO control (Fig. 2E). This suggested that M47 binds the computationally predicted amino acids in the PHR domain.

We hypothesized that M47 binds to the PHR domain and increases the degradation rate of CRY1 by increasing its ubiquitination level ^28–30^. To test this HEK293T cells were transfected with pcDNA-Cry1-His-Myc, Fbxl3-GAL4, and pUb-HA plasmids in the presence and absence of M47. To determine the ubiquitination level of CRY1, proteasomal degradation was blocked by MG132. Proteins were isolated with anti-Myc resin and ubiquitination levels were determined by Western blot. The analysis of the ubiquitination assay revealed that CRY1 is hyper-ubiquitinated when treated with M47 compared to DMSO control (Fig. 2F).

All these results showed M47 binds to the FAD-binding pocket in the PHR domain and specifically destabilizes CRY1 by enhancing the ubiquitination of CRY1.

### Physical Interaction between M47 and CRY1

To show the physical binding of the M47 to CRY1, biotinylated-M47 (bM47) was commercially synthesized by (Enamine, Ukraine) (Fig. S3B). To test whether bM47 binds CRY1 plasmids have His-Myc tagged *Cry1* and *Cry2* were transfect to HEK293T cells. After preparation of the cell lysate, bM47 was used to pull-down CRY1-HIS-MYC and CRY2-HIS-MYC in the presence and absence of the competitor (free M47). Results indicated that M47 physically binds to CRY1 but not to CRY2 (Fig. 3A). The binding of bM47 to CRY1 disappeared in the presence of the competitor. To confirm the binding of the molecule to the PHR domain of the CRY1 we performed a pull down assay between bM47 and CRY1-T5 (the lack of 100 amino acids from the C-terminal end). The result showed M47 bound to the PHR domain (Fig. 3B). The activity of biotinylated M47 was confirmed on circadian rhythm and half-life of CRY1 to make sure that biotinylation did not alter the function and binding of the molecule (Fig. S3C, D). Pull-down assays along with mutagenesis studies suggested that M47 physically binds to CRY1 through the FAD-binding pocket.

**Fig. 3.**
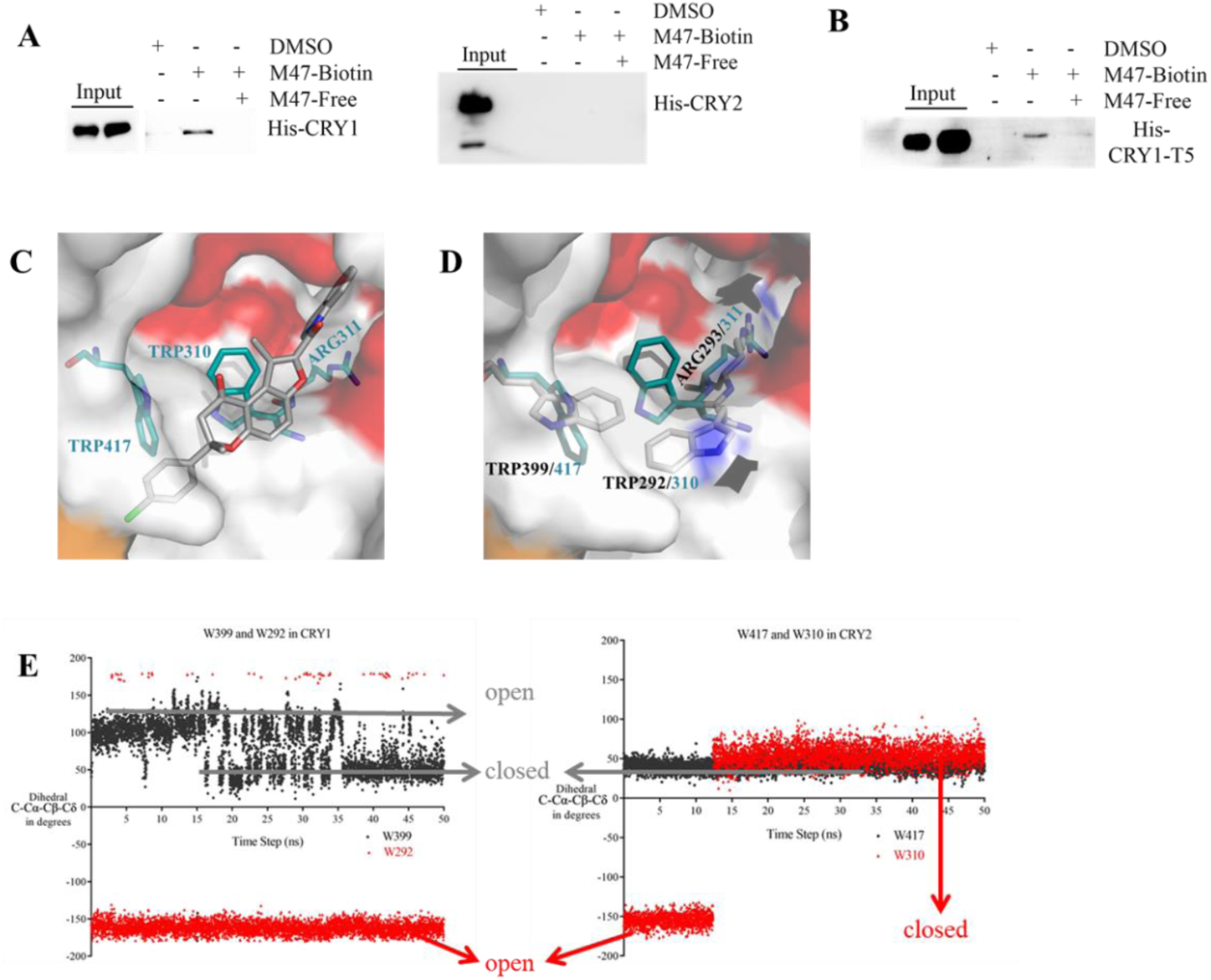
Physical interaction between the M47 and CRYs. (**A,B**) HEK293T cells transfected with *Cry1-His-Myc*, *Cry2-His-Myc*, or *Cry1-T5-His-Myc* plasmids then lysates were subjected to pull-down assay. Lysates treated with solvent (DMSO), 50µM bM47, and 50µM bM47 with 100µM M47 (competitor). While M47 binds to PHR of CRY1 (A, B) it does not bind to CRY2 (n=3). (**C**) Binding mode of M47 on equilibrated CRY2 (4I6G) predicted by AutodockVina with binding energy −9.4kcal/mol. W310 occupies the binding region of M47. Coloring is done as explained in Fig. 2D with modifications. Carbon atoms of CRY2 are shown in cyan. (**D**) Superimpose image of equilibrated CRY1 and CRY2 structures. M47 binding residues (W399, W292, R293) in CRY1 and corresponding residues in CRY2 (W417, W310, R311) are shown in sticks. (**E**) Dihedral angle between C-Cα-Cβ-Cδ atoms of W399 and W292 in CRY1; W417 and W310 in CRY2 throughout the simulations. Dihedral angle around −150° of W292 in CRY1, W310 in CRY2 represents the parallel position; angle > 0° represents the perpendicular position of these residues to R293 and R311 in CRY1 and CRY2, respectively. Similarly, dihedral angle around 90° of W399 in CRY1, W417 in CRY2 represents the parallel position; angle around 40° represents the perpendicular position of these residues.

To understand why CRY2 was unable to bind M47 we performed computational studies on the FAD-binding region of the CRY2. Similar to CRY1, we docked M47 to CRY2. The docking position of M47 on CRY2 showed that the molecule can not interact with W417 and R311 of CRY2, which are homologous to W399 and R293 of CRY1 (Fig. 3C). Autodock Vina predicted the binding energy of M47 to CRY2 and CRY1 as −9.5 and −13.3 kcal/mol, respectively. To further understand the difference in the binding residues of M47 between CRY1 and CRY2, we analyzed the equilibrated CRY1 and CRY2 structures used in docking simulations. Although residues in PHR of CRY1 and CRY2 are 77% identical and 88% similar, two of the critical residues (W399 in CRY1, W417 in CRY2; W292 in CRY1, W310 in CRY2) locate themselves differently in equilibrated CRY1 and CRY2 structures (Fig. 3D). To understand the conformational preferences of amino acid residues interacting with M47 in both CRYs in detail, dihedral of C-Cα-Cβ-Cδ atoms were measured throughout the 50ns MD simulations. Dihedral angle around 50° for W399 CRY1 (W417 in CRY2) corresponds to conformation where tryptophan residue is perpendicular to R293 in CRY1 (R311 in CRY2) and the pocket is closed for molecule binding, where values around 100° correspond to the parallel position of these two residues and the pocket is open for molecule binding. Dihedral value around −150° for W292 in CRY1 (W310 in CRY2) corresponds to the parallel position of this residue to R293 in CRY1 (R311 in CRY2) (open), however, >0° corresponds to perpendicular position (closed) (Fig. 3E). While side chains of W399 and W292 were mostly located parallel to R293 in CRY1 and left enough space M47 to bind, W417 and W310 in CRY2 acquired a position perpendicular to R311 where they occupied most of the M47 binding space in CRY2. Although W399 mainly maintained its position parallel to R293 in CRY1 (76% of the entire simulation), W310 kept its position perpendicular to R311 in CRY2 and filled the binding space of M47 through the entire simulation. Superimposing the equilibrated structures of CRY1 and CRY2 showed how these critical amino acid residues were located differently (Fig. 3E). These results showed that the internal dynamics of CRY1 and CRY2 could be different even at the very conserved region which can modulate their interactions with proteins or ligands, and thus their cellular roles.

### Time-Dependent Effect of M47 on the U2OS Cells

To evaluate the effect of M47 on the endogenous protein and mRNA level of clock genes, synchronized U2OS cells were treated with 10 µM M47. Cells were harvested between 24-48 h of post-synchronization at 4 h time intervals. M47 decreased the level of CRY1, all the time points but 36^th^ h (Fig. 4A). On the other hand, we observed no significant change in CRY2 and CLOCK levels. There was a slight change in the BMAL1 level. Analysis of mRNA levels of *Cry1* and *Cry2* showed that M47 did not change their overall abundance (Fig. 4B). An increase in clock output genes of the *Dbp* and *Per2* mRNA levels also confirmed the decrease in the total repressor capacity of the cells treated with M47. Observing the effect of M47 on CRY1 at the protein level but not at the mRNA level confirms the post-translational effect of the molecule. All these results suggest that M47 is a promising molecule to regulate circadian clock machinery through CRY1. To investigate pharmacological properties and activity of M47 *in vivo*, it has been subjected to in mice studies.

**Fig. 4.**
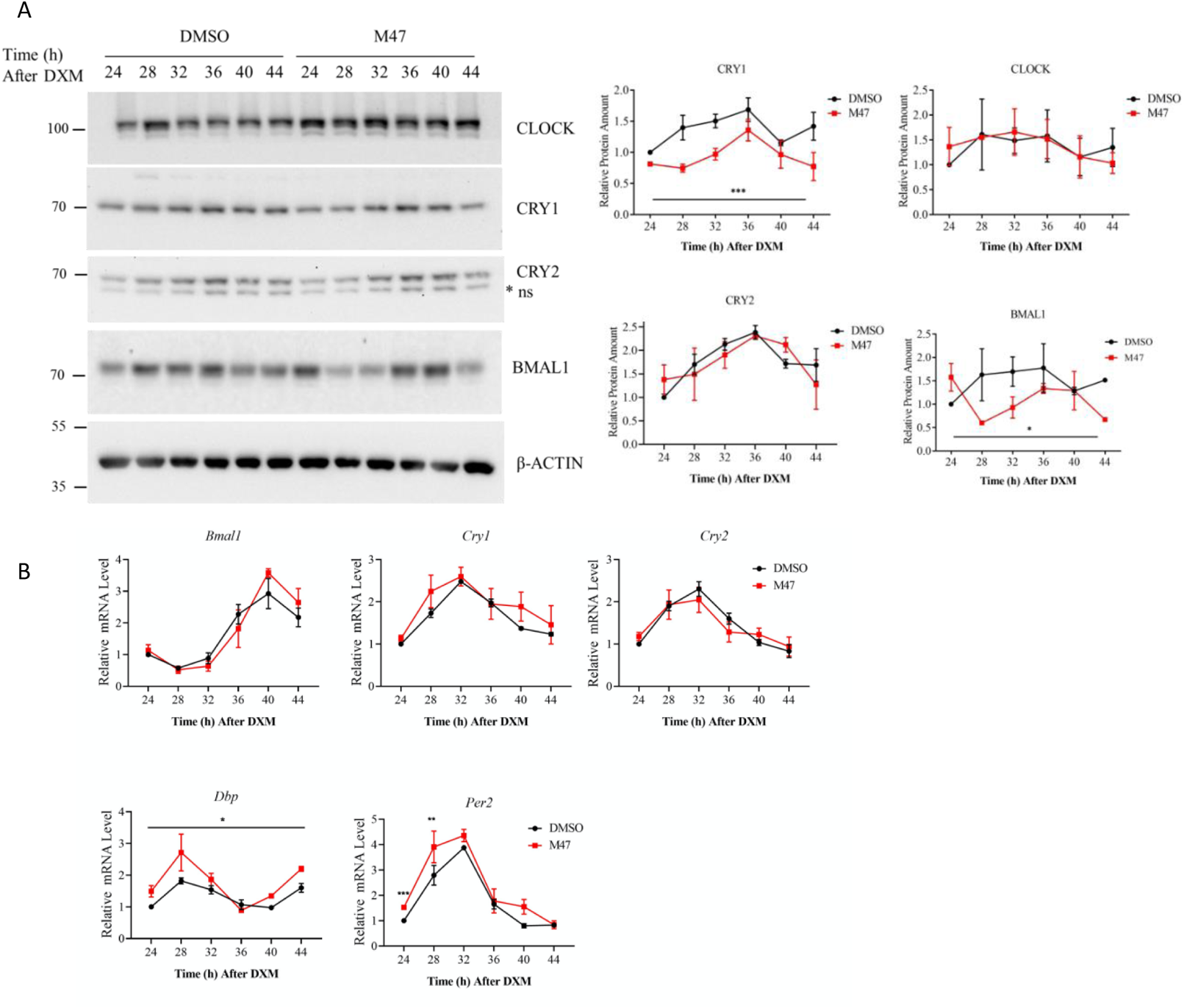
M47 decreased CRY1 level in synchronized U2OS cells. Confluent U2OS cells were synchronized by 2h treatment with dexamethasone (0.1µM) and medium replaced with fresh medium containing M47 or solvent (DMSO final 0.5%). Cells were harvested at indicated time points. (**A**) Lysed cells were analyzed via protein immunoblot technique. M47 decreased the protein level of CRY1 between 24^th^-48^th^ h. The level of proteins were normalized according to DMSO at 24^th^ h (mean ± SEM, n=3). (**B**) Cells were subjected to reverse-transcription-quantitative polymerase chain reaction (RT-qPCR). M47 treatment increased overall *Dbp* level and not affect the *Cry1* and *Cry2* levels (*: p < 0.05 versus DMSO control by two-way ANOVA). At 24 and 28^th^ hours M47 increased *Per2* (student’s t-test **: p<0.01, ***:p<0.001) Expression level of genes was normalized according to the expression level of that gene at 24^th^ h. (Data represent the mean ± SEM, n=3 with duplicates) *ns=non-specific band in western blot.

### Toxicity Studies in Mice

To evaluate the toxicity of the M47 *in vivo* we first performed a single-dose toxicity study (SDT). M47 was intraperitoneally (i.p.) administered to C57BL/6J mice at the doses of 5, 50, 300, and 1000 mg/kg. General toxicity was evaluated based on mortality, body weight changes, body temperature, clinical signs, food and water consumption, behavior assessment, and gross findings at terminal necropsy. Animals treated with 1000 mg/kg of the M47 exhibited the following clinical symptoms: dyspnea, hyporeflexia, excessive hypothermia, reduced locomotor activity, piloerection, hunched posture, and corneal opacity. This dose was considered lethal, and animals were sacrificed for ethical reasons. No mortality and clinical signs were observed in other doses (5, 50, 300 mg/kg). Compared to control mice (treated with the vehicle where their body temperatures were 36.0-37.0 ͦ C) mild hypothermia (33.5-35.1 ͦ C) was observed in mice within 6h after administration of the M47 and body temperatures gradually increased in the next hat the dose of 300 mg/kg (Fig. 5A). The body temperature of the animals was comparable to control groups at the dose of 5 mg/kg while the body temperatures of the animals were higher in animals treated with 50 mg/kg (Fig.5A). We then measured weight loss on these animals for 15 days. The highest mean body weight loss (7.7%) occurred on the first day following 300 mg/kg M47 administration (Fig 5B). There were no significant weight losses in animals treated with 5 mg/kg of the M47. All these results suggested that STD of 5 and 50 mg/kg were well-tolerated and 300 mg/kg tolerated by mice with some adverse effects. We next determined the 5-day maximum tolerated dose (MTD) using 40, 80, or 150 mg/kg of i.p. M47 with repeated injections for 5 days. A single death (1/6; 16.6%) among all tested doses was observed in the group of 150 mg/kg on the 3^rd^ day of injection. There were no observed clinical signs for the animals treated with 40 and 80 mg/kg of the M47. Hyporeflexia was observed in animals treated with 150 mg/kg of the M47, which was more prominent in the first three days of treatment and decreased on the 4^th^ and 5^th^ days. The body temperature of the animals treated with 40 mg/kg of M47 was comparable to the control animals. On the other hand, mild hypothermia (33.4-35.8 ͦ C) was measured in mice on the first day at doses of 80 and 150 mg/kg. Body temperatures increased gradually in the following days (Fig. 5C). The body weight losses were similar for all doses with an exception at 3^rd^ and 4^th^ injections at the dose of 150 mg/kg where the highest mean body weight loss (6.3%) was observed on the 3^rd^ day of injection (Fig. 5D).

**Fig. 5.**
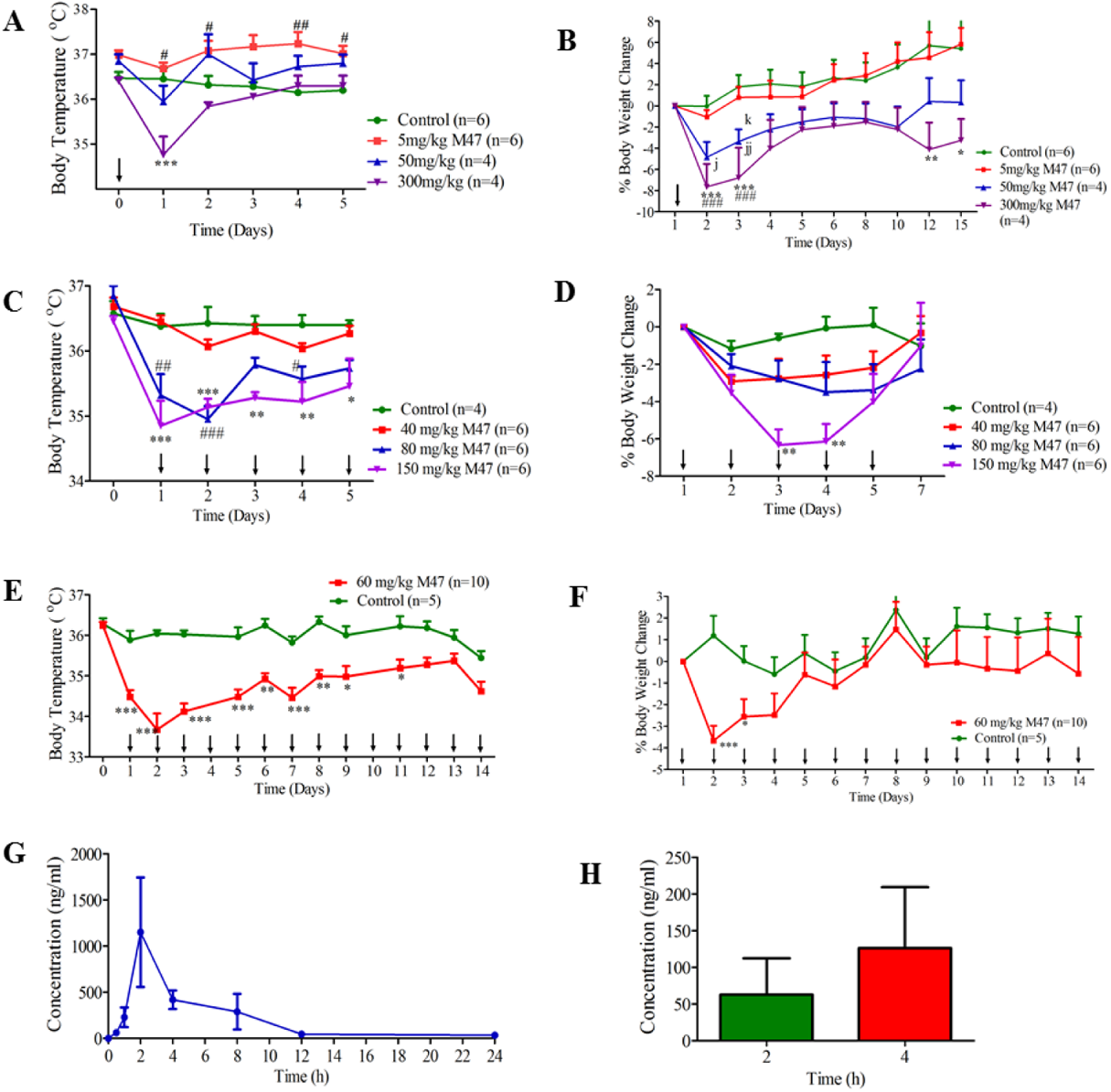
Toxicity and pharmacokinetics of M47 in mice. (**A**) Body temperatures in C57BL/6J mice treated with single intraperitoneal doses of 5 mg/kg, 50 mg/kg, 300 mg/kg M47 or vehicle during 15-day observation period. Body temperatures (ͦ C) were expressed as mean ± SEM. “↓” indicates the treatment days of M47. ***p<0.001, control vs 300 mg/kg M47; ^#^p<0.05, ^##^p<0.01 control vs 5 mg/kg M47 (Two-way ANOVA with Bonferroni post hoc test). (**B**) Body weight changes (%) in C57BL/6J mice treated with single intraperitoneal doses of 5 mg/kg, 50 mg/kg, 300 mg/kg M47 or vehicle during 15-day observation period. Body weight changes (%) were expressed as mean ± SEM. “↓” indicates the treatment days of M47. *p<0.05,**p<0.01, ***p<0.001, control vs 300 mg/kg M47; ^#^p<0.05, ^##^p<0.01, ^###^p<0.001 5 mg/kg vs 300 mg/kg M47; ^j^p<0.05, ^jj^p<0.01 control vs 50 mg/kg; ^k^p<0.05, 5 mg/kg vs 50 mg/kg M47 (Two-way ANOVA with Bonferroni post hoc test). (**C**) Body temperatures in C57BL/6J mice treated with intraperitoneal doses of 40 mg/kg, 80 mg/kg, 150 mg/kg M47 or vehicle for 5 days. Body temperatures ( ͦ C) were expressed as mean ± SEM. “↓” indicates the treatment days of M47. *p<0.05,**p<0.01, ***p<0.001, control vs 150 mg/kg M47; ^#^p<0.05, ^##^p<0.01, ^###^p<0.001 control vs 80 mg/kg M47 (Two-way ANOVA with Bonferroni post hoc test). (**D**) Body weight changes (%) in C57BL/6J mice treated with intraperitoneal doses of 40 mg/kg, 80 mg/kg, 150 mg/kg M47 or vehicle for 5 days. Body weight changes (%) were expressed as mean ± SEM. “↓” indicates the treatment days of M47. **p<0.01 control vs 150 mg/kg M47 (Two-way ANOVA with Bonferroni post hoc test). (**E**) Body temperatures in C57BL/6J mice treated with intraperitoneal dose 60 mg/kg M47 or vehicle for 14 days. Body temperatures ( C) were expressed as mean ± SEM. “↓” indicates the treatment days of M47. *p<0.05, **p<0.01, ***p<0.001, control vs 60 mg/kg M47 (Two-way ANOVA with Bonferroni post hoc test). (**F**) Body weight changes (%) in C57BL/6J mice treated with intraperitoneal dose of 60 mg/kg M47 or vehicle for 14 days. Body weight changes (%) were expressed as mean ± SEM. “↓” indicates the treatment days of M47. *p<0.05, ***p<0.001, control vs 60 mg/kg M47 (Two-way ANOVA with Bonferroni post hoc test). (**G**) Mean plasma concentration-time curve of M47 administered at 100 mg/kg intraperitoneally to C57BL/6J mice. Data were expressed as mean ± SEM (n=4 per time point). (**H**) M47 levels in brain tissue at 2^nd^ (n=2) and 4^th^ (n=2) hours following M47 administration at a dose of 100 mg/kg intraperitoneally to C57BL/6J mice. Data were expressed as mean ± SEM.

All these results suggested that doses of 5 and 80 mg/kg were well-tolerated by mice. We, therefore, decided to carry out a subacute toxicity test to further evaluate the effect of the M47 on animals using the dose of 60 mg/kg for 14-days of repeated injections. When M47 was administered to animals once a day for 14 days no mortality and clinical signs were observed. Mild hypothermia (33.7-35.4 ͦ C) was recorded in mice during injections (Fig. 5E). The highest mean body weight loss (3.7%) occurred on the first day following administration and body weights increased through the next 13 days of injections (Fig. 5F).

### Determination of Pharmacokinetic Profile of M47

The mean plasma concentration-time curve of M47 administered at 100 mg/kg single dose (i.p.) to female C57BL/6J mice were presented in Fig 4G, and pharmacokinetic parameters of M47 were given in Table 1. M47 plasma level was first detected at 0.5 h, reached a maximum level at 2-4h and was still quantifiable at 24 h. For the assessment of the brain exposure of M47, levels in this tissue were only determined at 2^nd^ (n=2) and 4^th^ (n=2) haccording to maximum concentrations of the molecule in plasma. M47 was detected in the brain tissue (Fig. 5H). All these results suggested that the M47 has a half-life *in vivo* and crosses the blood-brain barrier.

**Table 1.** Plasma pharmacokinetic parameters of M47 (100 mg/kg, single dose, i.p.)

| **Parameters** | **Values** |
| --- | --- |
| C<sub>max</sub> (ng/ml) | 1150.52±506.26 |
| T<sub>max</sub> (h) | 2-4 |
| AUC<sub>0-24</sub> (ng.h/ml) | 4921±3069.09 |
| AUC<sub>0-∞</sub> (ng.h/ml) | 5194±3134.00 |
| k<sub>el</sub> (1/h) | 0.127±0.01 |
| t<sub>1/2</sub> (h) | 5.5±0.33 |

### *In vivo* effects of M47

All *in vitro* studies suggested that M47 binds and reduces the half-life of CRY1. To assess its *in vivo* effect, M47 was i.p. administered to mouse at 25mg/kg and 50 mg/kg doses. Mice were sacrificed at 6^th^ h after the injection and all organs were kept for downstream analysis. We observed a slight decrease in CRY1 levels in the whole cell lysate of mouse liver cells (Fig 6A). Since CRYs are stabilized and degraded differently in cytosolic and nuclear compartments ^9, 10^, we further analyzed the level of CRY1 in the cytosolic and nuclear fraction of the mouse liver. Proteins, known to be specifically localized in nucleus (Histone-H3) and cytoplasm (Tubulin), were used as controls to evaluate the purity of the fractions (Fig. S4). CRY1 abundance in the nucleus was significantly low as compared to controls in dose-dependent manner (Fig. 6B). On the other hand, cytosolic levels of CRY1 in M47 treated mice liver were not statistically significant as compared to controls (Fig. 6C). To analyze the effect of M47 on the clock-dependent transcription, total RNAs were isolated from the same samples and mRNA levels of *Per2* were measured by qPCR. In agreement with decreased CRY1 level in the nucleus, the expression level of *Per2* significantly increased in mice treated with M47 (Fig. 6D). All these suggested that M47 is effective on CRY1 levels in mouse liver cells.

**Fig. 6.**
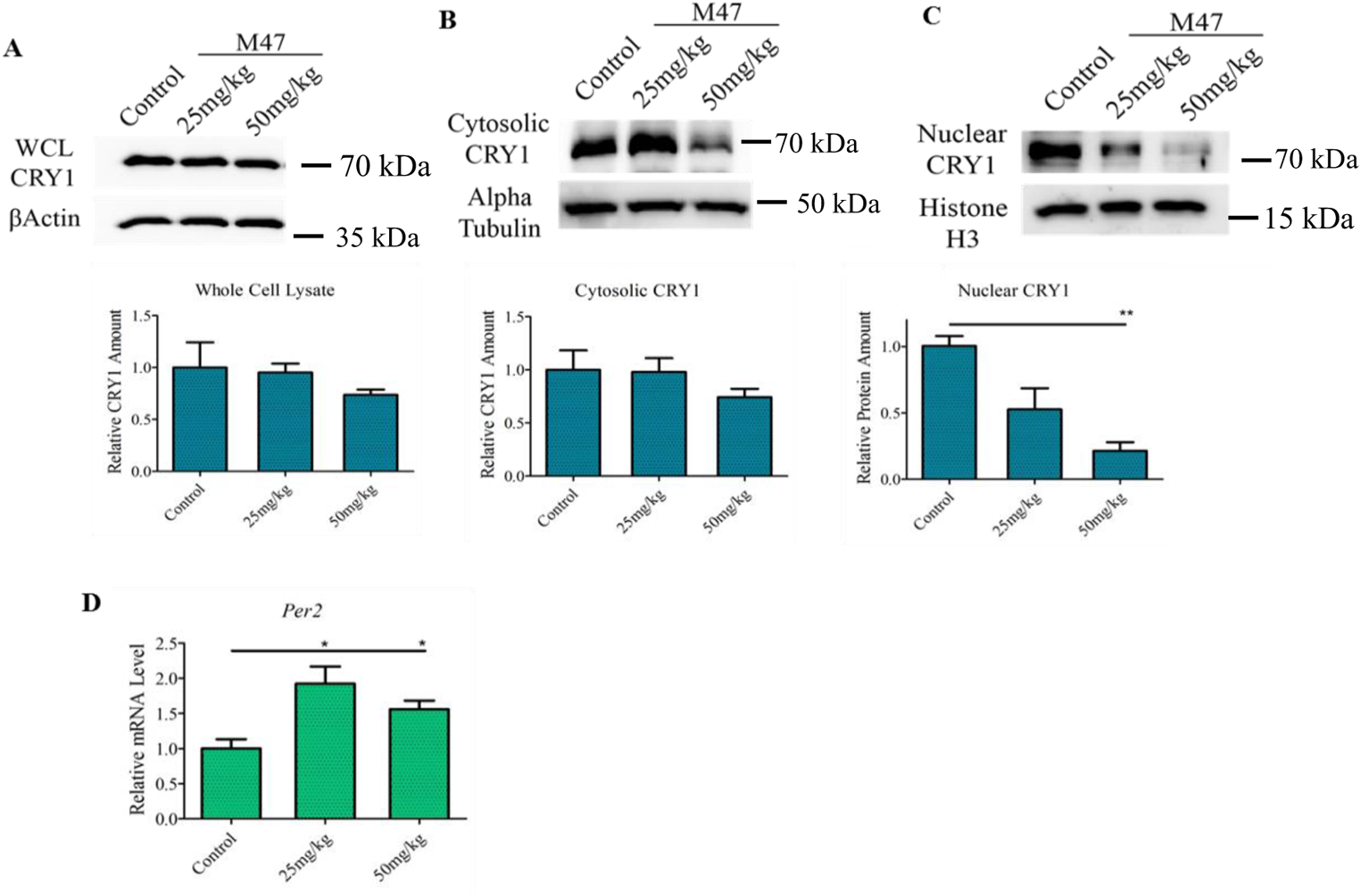
M47 decreased the nuclear CRY1 in mice liver. M47 (25 mg/kg or 50mg/kg) or vehicle intraperitoneally administered to C57BL/6J mice. 6h after the treated mice were sacrificed (n=3). While M47 slightly decreased the CRY1 levels in whole cell lysate (WCL) and cytosolic fraction, decreased the nuclear CRY1 level significantly. Effect of M47 in (**A**) WCL cytosolic fraction, and (**C**) nuclear fraction of mice liver. (**D**) Liver samples subjected to RT-PCR. M47 treatment increased the mRNA level of *Per2*. (Data represent the mean ± SEM, n=3 (with duplicates in RT-PCR) *: p < 0.05 versus control by student’s t-test).

### Effect of M47 on Apoptosis in p53^−/−^ Mouse Embryonic Fibroblast Cells

A comparative study showed that treatment of fibroblasts isolated from the skin of *Cry1^−/−^ Cry2^−/−^p53^−/−^* and *p53^−/−^* mice with UV or UV mimetic agents e.g oxaliplatin sensitizes the cells to bulky-DNA adduct-induced apoptosis when *Cry* deleted on the p53-null background ^31^. With the expectation that M47 would destabilize CRY1 and increase the effectiveness of UV mimetic chemotherapeutic agents, we performed an apoptosis assay with a Ras-transformed p53-null MSF cell lines. In this assay, cells were treated by increasing dose of oxaliplatin for 16h followed by M47 treatment for 24h and probed for apoptosis by measuring PARP cleavage. As a chemotherapeutic agent, oxaliplatin was preferred since mutation of *Cry* sensitizes p53-null cells to apoptosis ^32^. We found that M47 treatment sensitized a Ras-transformed the p53-null MSF cell line to oxaliplatin-induced apoptosis in dose-dependent manner (Fig. 7A). In the same setup, we checked the level of CRY1 and verified that the CRY1 levels were reduced in MSF cells when treated with M47.

**Fig. 7.**
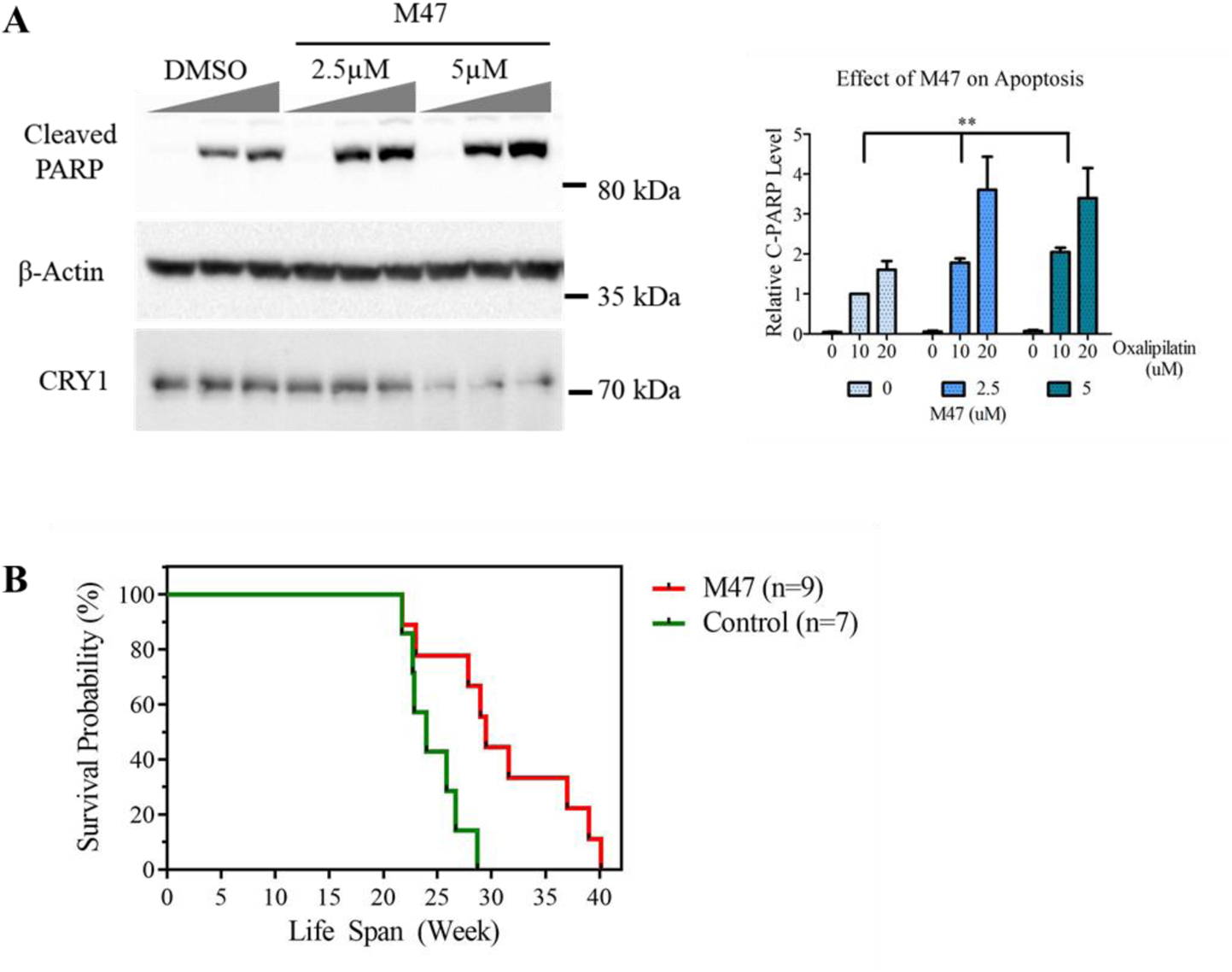
M47 enhanced the effect of oxaliplatin in *p53* null mouse skin fibroblast (MSF) cells. (**A**) Ras pT24 transformed p53 null MEF fibroblast cells were treated with 0, 10 or 20µM oxaliplatin and incubated for 24h. Then either DMSO or M47 was added and incubated for 16h. Cells were lysed and analyzed via protein immunoblot technique. At each cases M47 increases the cleaved PARP protein level and decreases CRY1 levels. Bar graph was drawn normalizing to 10µM dosage of oxaliplatin (Data represent the mean ± SEM, n=4 **: p < 0.005, versus DMSO control by two-way ANOVA). (**B**) M47 in *p53*^−/−^ mice reduces age adjusted tumor incidence. Kaplan–Meier survival analysis (log-rank test) of the time of death with evidence of tumors showed significant differences between vehicle treated *p53*^−/−^ and M47 treated *p53*^−/−^ (**p<0,01)

### Effect of M47 on the lifespan of p53^−/−^ mice

All our results showed that M47 selectively reduces the half-life of CRY1 and is well tolerated with a good pharmacokinetic profile in vivo and enhances the apoptosis. Studies showed that *p53*^−/−^ mice mostly develop lymphomas and lymphoid sarcomas with an average of 20 weeks in C57BL/6J background ^33, 34^. It is well-established that cancers develop, mostly, with a median age of about 20 weeks in C57BL/6J background ^34^. Additionally, as previously shown, deletion of *Cry* genes in *p53^−/−^* in C57BL/6J background enhances the lifespan of the animals by protecting animals from cancer death ^35^. A recent transcriptomic study revealed that differential regulation of nuclear factor kappa B (NF-κB) regulation may increase survival rate ^36^. We wished to test M47 on *p53^−/−^* animals to see whether it delays cancer death and change the lifespan in cancer prone *p53^−/−^* in C57BL/6J mice. To test that 50 mg/kg/day M47 was intraperitoneally administered to the *p53^−/−^* mice for 5 days per week, for 7 consecutive weeks (n=9) along with vehicle-treated control animals (n=7). As seen in Fig. 7B, the age-adjusted incidence of cancer in M47 treated *p53^−/−^* mice was significantly higher than that of vehicle treat *p53^−/−^* mice. The median lifespan of M47 treated *p53^−/−^* mice was 29.5 weeks while the median lifespan of vehicle-treated *p53^−/−^* mice was 24 weeks. Our results suggest that reducing the level of CRY1 protects mice from tumor burden and increases the survival rate of *p53^−/−^* mice.

## Discussion

Given the increasing prevalence of circadian disruptions and its deleterious health consequences, it is important to prevent and minimize circadian disruptions in our daily life and their detrimental effects ^37^. The study of fast, dynamic biological processes demands the development of approaches that conditionally and selectively modulate a specific aspect of a gene product in a rapid and tunable manner. In addition, all information regarding how clockwork is regulated at the molecular level comes from genetics studies. Despite the specificity and robustness of genetic manipulation, the lack of temporal control makes genetic approaches problematic for the study of biological processes occurring on the millisecond to minute time scale (*e.g.*, posttranslational modification, signal transduction) ^38^. Treatment strategies involving genetic or alterations in a patient’s daily behavior are not ideal for many reasons. A more promising therapeutic approach would be to use a pharmaceutical agent that selectively targets the molecular clock. These challenges will be overcome by discovering the small molecules that specifically bind and regulate the activity of each of these core clock proteins. Currently, a growing number of groups are looking for interventions ranging from modulating environmental cues to manipulation of molecular machinery. Therefore, molecules that either bind to modifiers of core clock proteins, such as casein kinase Iε (CK1ε), glycogen synthase kinase 3β (GSK3β), AMP-activated protein kinase (AMPK), and SIRT1 ^39–41^ with varying effect on the speed of the clock and amplitude ^42^; or that target CRY ^17, 25, 43^, REV-ERBs ^18, 44^, and RORs ^45, 46^, three of core clock proteins, were identified from high-throughput screening.

For the first time structure-based drug design approach unveiled a CRY1 binding molecule, M47, which decreases the half-life and overall CRY1. Our findings demonstrated that M47 binds to CRY1 but not CRY2, reduces the half-life of CRY1-LUC when overexpressed in HEK293T cells and decreases overall CRY1 abundance both in U2OS cells and mouse liver cells. In synchronized U2OS cells M47 decreases CRY1 level at certain time points and increases circadian clock output genes, e.g. *Per2* and *Dbp* as expected when repressor capacity is attenuated. Docking simulations suggested that M47 binds to the PHR region of CRY1 which is a critical domain for the rhythmicity of the cells ^45^. Docking pose and mutational analysis show that M47 binds to W399 and R293 in the FAD-binding pocket. Despite a decrease in the level of CRY1, M47 lengthens the period and dampens the amplitude in U2OS *Bmal1*-d*Luc* cells. One possible explanation for the unexpected period lengthening is that since R293 is located just behind the secondary pocket, the interaction between M47 and R293 may stabilize CRY1-CLOCK complex which was shown to be critical for period length and repression activity ^47, 48^. Cell-based genetic complementation studies also show that the stability and amount of CRY, *per se*, do not dictate the period length, but that the subcellular localization and interaction of CRY with other core clock proteins determine the period ^5, 16, 49–51^. Although period shortening molecules were previously expected to be CRY destabilizers, GO044, GO200, and GO211 shortened the period while enhancing the CRY stability ^25^. This strongly supports the complicated relation between CRY half-life and the period length.

Another distinct property of M47 is its preferential binding to PHR of CRY1 but not CRY2 which are 77% identical. Although binding residues of M47 (R293, W292 and W399 in CRY1; R311, W310, and W417 in CRY2) are conserved between CRY1 and CRY2, detailed analysis of MD simulations under physiological conditions showed that even these conserved residues prefer different conformations. Different conformational preferences allowed M47 to specifically bind to CRY1 but not CRY2 due to the steric hindrance governed by W417 and W310 in CRY2. This difference implies that CRY1 and CRY2 may have diverse dynamics despite their high similarity. Our results suggest that differences between CRY1 and CRY2 dynamics can be utilized to develop small molecules specifically regulating the activity of each protein which would contribute to a better understanding of the roles of CRY1 and CRY2 in the clock-work mechanism.

Pharmacological studies revealed that animals tolerated well single and repeated subacute doses of M47 with a half-life of around 6h in blood. M47 also crosses the blood-brain barrier without altering the hematological parameters. All these results suggest that M47 may be a drug-like molecule. Further *in vivo* studies indicated that M47 also interacts with CRY1 and reduces its level in the mouse liver cells.

Importantly, knockdown of the *Cry*s in *p53^−/−^* mice or cell lines improved sensitivity to cancer chemotherapy by activating tumor suppressor genes ^14, 52^. In addition, overexpression of *Per2* genes in human pancreatic cancer cells prevented cell proliferation, initiated apoptosis and behaved synergistically with cisplatin ^53^ thereby suppressed the tumor growth. Studies on the role of *Cry* and *Per2* in cancer suggested that CRY destabilizers, expected to be P*er2* enhancers, can be utilized as novel anti-cancer therapeutics. Since M47 decreases CRY1 and enhances the level of *Per2* both in cell-line and *in vivo,* we hypothesized that M47 could be a useful molecule for cancer by promoting apoptosis. Thus, we analyzed the effect of M47 on the Ras-transformed *p53*-null fibroblast cells treated with oxaliplatin used as UV-mimetic. M47 enhanced the effect of oxaliplatin which implies that M47 can be utilized as an anti-cancer therapeutic to increase the effect of oxaliplatin. Finally, we showed that administrating the M47 to *p53^−/−^* mice increased their life span ~25%. This result is consistent with the previous study, where knocking out *Cry* genes in *p53* null mutant improved the lifespan of *Cry* mutant mice ^14^.Collectively, our data suggest M47 has great potential to be used as a chemotherapeutic molecule for cancer types having p53 mutations.

## Materials and Methods

### Molecular Dynamics Simulation

Mouse CRY1 (PDB ID: 4K0R) and mouse CRY2 (PDB ID: 4I6G) structures were obtained from protein databank. For each structure, using the NAMD (v. 2.6) and VMD (v. 1.9.1) program packages each protein was solvated in a rectangular box with TIP3P water molecules and neutralized with counter ions as describe in ^54^. Then the system was minimized and heated up to physiological temperature with 10K increments by running 10ps simulation at each temperature. CHARMM-PARAM22 force field was used for the molecular dynamics (MD) simulations. After the equilibration of the system, MD simulation was run at 310°K for 50 ns. Pressure was controlled by Langevin piston method during the simulations. Time step was set to 2fs and the bonded interactions, the van der Waals interactions (12Å cutoff), long-range electrostatic interactions with particle-mesh Ewald (PME) were included for calculating the total force acting on the system. Equilibrated CRY1 structure was used as the “receptor” for the docking simulations. The root-mean square deviation (RMSD) calculations were done using VMD RMSD utilities. Backbone atoms (C, CA, and N) of each residue were used for RMSD calculation by excluding their translational motions.

### Molecular Docking

More than 8 million small molecules, freely available from Ambinter Web site, with non-identified functions were used as ligands for the docking. Molecules were filtered according to the following criteria to eliminate non-relevant molecules: molecules should have less than 7 H-bond donor, less than 12 H-bond acceptor, less than 600 Da molecular weight, logP < 7, less than 8 rotatable bonds, at least 3 aromatic rings, and 04 at least 4 rings. Openbabel, Autodock4.2, Autodock Tools4 ^55^ and Autodock Vina ^56^ program packages which are free for academic purposes, were utilized to prepare ligands (small molecules) for the docking. Finally, more than 1million compounds were docked to target pockets by using the Autodock Vina program. Target pocket for FAD and FBXL3 binding site was determined based on the CRY-FBXL3 crystal structure ^57^. Autodock Tools4 or PyMol (http://pymol.sourceforge.net/) software were used to visualize the docking results and protein structure, respectively.

### MTT-Toxicity Assay

Human osteosarcoma U2OS and NIH 3T3 cell lines were used for the cytotoxicity assay. Cells were seeded in triplicate to clear 96-well plates with 4000 cells/well then grown for 48h. Cells were treated with molecules at desired concentrations (final DMSO concentration 0.5%) in DMEM. Cells were incubated with molecules for 48h. Cell viability was measured by adding tetrazolium dye 3-[4,5-dimethylthiazol-2-yl]-2,5 diphenyl tetrazolium bromide (MTT) which is converted to insoluble purple color formazan because of the mitochondrial activity. To measure the purple color after incubating cells with MTT reagent medium was replaced with DMSO:EtOH (50:50) mixture for 4 hand absorbance of wells were measured at 570nm by the spectrophotometer. Cells treated with 5% final DMSO concentration (known as toxic to cells) as negative controls.

### Real-Time Bioluminescence Monitoring

This assay was performed with two different equipment: Synergy H1 (BioTek) with 96-well plate and LumiCycle luminometer (Actimetrics) with 35mm plates. Initial screening of molecules to determine their effect on the circadian period was performed in 96-well plate via Synergy H1. Clear 35-mm plates are used in LumiCycle luminometer which provides high resolution data describing the circadian oscillation. For 96-plate tests, 50000 U2OS *Bmal1-dLuc* cells were seeded on an opaque 96-well plate and cultured overnight. Next day cells were reset by adding dexamethasone (DXM) (0.1µM final) for 2h. Then medium is changed to bioluminescence recording media which contains the following in 1L: DMEM powder (sigma D-2902, 10X 1L), 0.35gr sodium bi-carbonate (tissue culture grade, sigma S5761), 3.5gr D(+) glucose powder (tissue culture grade, sigma G7021), 10mL 1M HEPES buffer (Gibco 15140-122), 2.5 mL Pen/Strep (100ug/ml), 50mL 5% FBS and up to 1L sterile milliQ water. Luciferin is added freshly with 0.1mM final concentration. Molecules were added to the bioluminescence recording media at desired concentration (0.5% final DMSO concentration). Plates were sealed with optically clear film to prevent evaporation and gas exchange thereby to maintain homeostasis of the cells.

Luminescence values were recorded at 32°C for each 30 min with 15 Sec integration time via Synergy H1 luminometer for a week. For LumiCycle, 400×10^3^ U2OS *Bmal1-dLuc* or NIH3T3 *mPer2-dLuc* cells were seeded to 35mm plates and then procedure given above was followed with a change in the last step as described earlier ^58^. Plates were sealed with vacuum grease and placed to LumiCycle. Each plate was recorded continuously every 10 min for 70 Sec at 37°C via photomultiplier tubes. Period and amplitude data were obtained from LumiCycle Analysis software. To detect the effect of M47 in the absence of CRYs, *Cry1^−/−^/Cry2^−/−^* mouse embryonic fibroblasts (CRY-DKO MEFs) transiently transfected with pGL3-Per2-Luc (luciferase reporter) were used. 3×10^5^ CRY-DKO cells were seeded in 35mm clear plates. Next day, cells were transfected with 4000ng pGL3-Per2-dLuc via Fugene6 transfection reagent according to the manufacturer’s instruction. In short, 3:1 ratio of Fugene6 in µl against transfected DNA amount in µg was kept in transfections. 72 h after transfection cells were synchronized with DXM for two hours. Then, medium was changed with lumicycle medium having DMSO or molecule, sealed with vacuum grease, and placed to LumiCycle.

### CRY-LUC Cloning

To insert *Cry1-Luc*, *Cry2-Luc* constructs and only *Luc* into pcDNA4A-myc-his plasmid, coding sequence of mouse *Cry1* and firefly Luciferase in pG5luc plasmid (Addgene) was amplified with primers having EcoRV-NotI and NotI-XhoI flanking sites for *Cry*s and Luc, respectively (Table S1). Product size was verified by visualizing in agarose gel and product was isolated from the gel by using NucleoSpin PCR and Gel purification kit (Macherey Nagel). pcDNA4A plasmid having *Luc* was double digested with NotI and XhoI and used as insert; *Cry1* pcDNA4A-myc-his and *Cry2* pcDNA4A-myc-his plasmids were double digested with NotI and XhoI FastDigest enzymes (Thermo Scientific) and used as host plasmids. Hosts were treated with FASTAP (Thermo Scientific) to prevent self-annealing. After gel isolation and cleaning inserts with Gel purification kit (Macherey Nagel) and destination vectors were ligated by using T4 DNA ligase (Thermo Scientific).

### Site Directed Mutagenesis

Quick-change method was used for the site-directed mutagenesis as described in ^59^. The primers were designed to have around 30 base pairs with the designated base changed in the middle of the primer. All the primer sequences for site-directed mutagenesis can be found in Table S1. The PCR reaction mixtures contained 0.3 mM dNTP, 5 µl of 10X Phusion GC Buffer (Thermo Scientific), 3% DMSO, 1 µM of each primer, 50 ng of the template plasmid (mCry1 in pcDNA4A) and 1 unit of Phusion DNA polymerase in a 50 µl of final volume. The reaction conditions were set to 98°C for 30 Sec, 55°C for 30 Sec and 68°C for 5 min, 18 cycles. PCR products were visualized in 1% agarose gel. The samples with correct sized bands were digested with 1 unit of DpnI FastDigest enzyme (Thermo Scientific) for 1 h at 37°C and then 5µl of sample was transformed to *E.coli* DH5α cells. Colonies were picked to culture and then plasmids were isolated via miniprep (Macherey Nagel). Sanger sequencing (Macrogen, Netherlands) was utilized to confirm the mutations.

### CRY-LUC Degradation Assay

*Cry1*-d*Luc* and *Cry2*-d*Luc* constructs were used as described in our previous study ^47^. 40ng of *Cry1*-*Luc*, *Cry2-Luc*, mutant *Cry1-Luc* plasmids or 5 ng *Luc* plasmid were reverse transfected to 4×10^4^ HEK293T cells on opaque 96-well plate with flat bottom via PEI transfection reagent. 24h after transfection, cells were treated with molecules or solvent (DMSO). 24h of post molecule treatment cells were treated with luciferin (0.4 mM final) and HEPES (10 mM final and pH=7.2). After 2 h, cycloheximide (20 µg/ml final) was added to wells to stop protein synthesis. Plate was sealed with optically clear film and placed to Synergy H1. Luminescence readings were recorded every 10min at 32°C for 24 h. Half-life of protein was calculated via one-phase exponential decay fitting function in GraphPad Prism5 software. For each molecule or control at least three replicates were done in each experiment.

### Protein immunoblots

Cells were lysed in RIPA buffer (50 mM Tris, 150 mM NaCl, 1% Triton-X, 0.1% SDS) with fresh Protease inhibitor cocktail (PIC). After centrifugation at 13000rpm for 15 min at 4°C, supernatant was used to determine the protein amount using Pierce Protein Assay (Thermo Scientific) according to the manufacturer’s instruction. Mice liver samples were homogenized and lysed with Dounce homogenizer and RIPA buffer. After homogenization samples were incubated 10 min on ice. After centrifugation 01 13000rpm 15 min at 4°C supernatant was used for the whole cell lysate. Subcellular fractionation of liver samples: Liver samples were homogenized with cytosolic lysis buffer (10mM HEPES pH 7.9, 10mM KCl, 0.1mM EDTA, 0.05% NP40, proteasome inhibitor) by pipetting 40 times with a cut tip. Then samples were incubated 10 min on ice and centrifuged for 3 min at 3000rpm at 4°C. Supernatant was saved in a new tube, and pellet was processed for the nuclear fraction. Supernatant was centrifuged at 13000 rpm for 3min at 4°C where new supernatant saved as the cytosolic fraction. Pellet was wash with cytosolic buffer and centrifuged 3 min at 3000rpm at 4°C and pellet was dissolved in nuclear lysis buffer (20 mM HEPES pH 7.9, 0.4 M NaCl, 1 mM EDTA, 10% glycerol, protease inhibitor), sonicated (60% power, 2 times 10 Sec x 3 cycles), and centrifuged 13000rpm for 5 min at 4°C. Supernatant is the nuclear fraction. After determining the protein amount, all lysates described above were supplemented with 4X Laemmli buffer (277.8 mM Tris-HCl pH: 6.8, 4.4% LDS, 44.4% (w/v) glycerol, 0.02% Bromophenol blue, (freshly added) 5% volume of beta Mercaptoethanol) and boiled at 95°C for 10 min. Lysates mixed with Laemmli buffer were resolved in SDS-PAGE and transferred to PVDF membrane (Millipore). Membrane was blocked with 5% milk solution in 0.15% TBS-Tween 20 for 1 h and then incubated overnight with the following primary antibodies where they necessary: CRY1 (Bethyl Catalog No. A302-614A), CRY2 (Bethyl Catalog No. A302-615A), PER2 (Bethyl Catalog No. A303-109A), BMAL1 (Santa Cruz, sc-365645), anti-Myc (Abcam, ab18185), anti-His (Santa Cruz SC-8036) BetaActin (Cell Signaling, 8H10D10), AlphaTubulin (Sigma, T9026), HistoneH3 (Abcam, ab1791). After washing membranes with 3 times for 5min with 0.15% TBS-Tween, membranes were incubated 1h with appropriate HRP conjugated secondary mouse (Santa Cruz SC-358920) or rabbit (Cell Signaling, 7074) antibodies. ECL buffer system (Advansta WesternBright) was used to visualize HRP chemiluminescence via BioRad ChemiDoc Touch visualizer.

### RNA Isolation and Real Time (RT) Quantitative PCR (qPCR)

QIAGEN RNeasy Mini Kit was used to isolate the mRNA. Protocol provided by the manufacturer was followed. On-column DNase treatment was performed with the DNase enzyme provided with the kit. Dounce homogenizer was used for the homogenization of the liver samples and followed protocol provided by the manufacturer to isolate RNA. Quantity of RNA was determined by Nanodrop2000 (Thermo Scientific). Quality of RNA was verified by running in ~0.5% agarose gel prepared with DEPC water and visualizing under UV light. ~400ng of each RNA sample were subjected to first strand cDNA synthesis with MMLV-reverse transcriptase (Thermo) as the following: RNAs were mixed with 2µl Oligo(dT)_23_, 1µl 10mMdNTP mix, and nuclease-free H_2_O to a final volume of 10µl (mix1). Mix1 was incubated at 65°C for 5 min to denature the possible secondary structures of RNA and primers which may prevent long cDNA synthesis then mixture was incubated at 4°C for 5 min. Mixture2 (mix2) containing 2 µl 10X reverse transcription buffer, 0.2µl Ribolock RNase inhibitor, 1µl Revert Aid Reverse Transciptase and 6.8µl nuclease-free H_2_O was added to mix1. Mixture of mix1 and mix2 (totally 20µl) was incubated at 42°C for 1 hour and then at 65°C for 20 min to inactive the enzymatic activity. Volume of each sample was completed to 100µl with nuclease-free H_2_O. For quantitative real time PCR (qRT-PCR) analysis cDNAs were further diluted 5-fold (1:5). mRNA expression levels were calculated by qRT-PCR with the SYBR Green. *Gapdh* gene was used as an internal control. A sample qRT-PCR reaction is the following: 8µl SYBR Green, 3µl cDNA (from 5-fold diluted solution), 1µl forward and reverse primer mix (0.5µM final), 8µl nuclease-free H_2_O. All reactions were performed in biological triplicates (each triplicate with two technical replicates), and the results were expressed relative to the transcript level of *Gapdh* in each sample using the 2^−ΔΔCT^method. qRT-PCR was run with the following cycling protocol: 95°C 10 Sec denaturation, 57 to 62°C for 20 Sec annealing (the degree is determined by primer type) and 72°C for 30 Sec of elongation, 35 cycles. List of primers used for the qRT-PCR can be found in Table S2 indicated as RT_GeneName.

### Pull-down Assay with Biotinylated Molecule

10µg of *Cry1*-pcDNA4, *Cry2*-pcDNA4 or *Cry1*-T5-pcDNA4 plasmids expressing His and Myc at the C-terminal of CRYs were transfected to HEK293T cell. After 48h of transfection cells were harvested and lysed in lysis buffer (50 mM Tris pH 7.4, 2 mM EDTA, 1 mM MgCl2, 0.2% NP-40 (v/v), 0.1% sodium deoxycholate (w/v), 1 mM sodium orthovanadate, 1 mM sodium fluoride, and protease inhibitor cocktail (Thermo Scientific). Lysates were incubated for 10 min on ice then mixed gently and incubated for another 10 min on ice. Then lysate was centrifuged for 10 min at 7000g at 4°C. 5% of the supernatant was kept for input analysis. Remaining supernatant was divided to three and mixed with 2x binding buffer (100 mM Tris-HCl pH 7.4, 300 mM NaCl, 0.2% NP-40 (v/v), 2mM sodium orthovanadate, 2 mM sodium fluoride, and protease inhibitor) with 1:1 ratio, incubated with rotation having either of these: a) DMSO, b) biotinylated molecule, c) biotinylated molecule and the competitor for 2h. During this period NeutrAvidin Agarose resin (Thermo Scientific) was equilibrated with lysis buffer by mixing for 5 min via rotation at 4°C three times. After each mix period, resins were centrifuged at 2500g at 4°C for 2 min and supernatant was discarded. After 2h of mixing, lysates were incubated with equilibrated resins for 2h to isolate the proteins interacting with the molecule. After 2h, resins were centrifuged (2500g, 2 min, 4°C) supernatants were discarded and washed with 1x binding buffer six times. Finally, proteins bound to resin were isolated by adding 50µl 4X Laemmli buffer and boiling at 95°C for 10 min. Pulled-down samples were analyzed via western blot.

### Ubiquitination Assay

1200ng of *mCry1*-His-myc, 300ng Gal4-*Fbxl3*, and 200ng of *hUb-Ha* were transfected to HEK293T cells via PEI transfection reagent on 6-well plate. Cells treated with DMSO or M47 after 28 hof transfection. 42 hof post transfection cells were treated with MG132 to block proteosome dependent degradation or equal volume of DMSO to negative control cells. 48 hafter transfection cells were harvested. CRY1 proteins were purified using Myc resin (EZview™ Red Anti-c-Myc Affinity Gel, SIGMA-ALDRICH Catalog No: E6654) according to the manufacturer’s instruction. In short, cells were lysed with RIPA buffer (50mM Tris, 150mM NaCl, 1% Triton-X, 0.1% SDS) and mixed with equilibrated myc-resins and agitated for 2 hr at +4°C. After washing three times with RIPA buffer, proteins were eluted with 4X Laemmli buffer (277.8mM Tris HCl pH: 6.8, 4.4% LDS, 44.4% (w/v) glycerol, 0.02% bromophenol blue, freshly added 5% beta Mercaptoethanol). Ubiquitination and expression levels of CRY1 was detected by Western blotting with anti-HA (Santa Cruz Biotechnology, sc7392) and anti-Myc (Abcam, ab18185) antibodies, respectively. Anti-Gal4(DBD) (Santa Cruz, sc-577) was used to detect Gal4-FBXL3 levels in the input.

### Generation of CRY1 knockout U2OS cell line

CRY1 gene were targeted in U2OS cell line using the LentiCRISPRv2 system ^47^. LentiCRISPRv2-CRY1 construct which was described previously ^60^. Briefly, the annealed oligos to target CRY1 (Sense: 5′ CACCGCCTTCAGGGCGGGGTTGTCG 3′; and Antisense: 5′ AAACCGACAACCCCGCCCTGAAGGC 3’) was inserted into BsmBI of LentiCRISPRv2 plasmid (Addgene #: 52961).

The lentivirus preparation, transduction of U2OS cells and selection of the knockout candidates with puromycin (at 0.5 mg/mL concentration) were performed following a previously published protocol ^60, 61^. Cell lysates of the knockout candidates were analyzed for CRY1, CRY2 and Actin by immunoblotting using the following antibodies: anti-CRY1 (A302-614A, Bethyl Labs Inc. Montgomery, TX., USA), anti-CRY2 (A302-615A, Bethyl Labs), and anti-Actin (CST-4967S, Cell Signaling Technology, Boston, MA, USA). CRY1 immunoblotting showed the absence of CRY1 protein while CRY2 immunoblotting showed that CRY2 was intact in this cell line. Actin immunoblotting was used as a loading control. HRP-labeled anti-mouse and anti-rabbit antibodies (Cell Signaling Technology) were used at 1:5000 dilution. Chemiluminescence was developed using WesternBright Sirius HRP substrate (Advansta, San Jose, CA, USA, cat no: K-12043-D20). Images of immunoblots were captured using the ChemiDoc XRS+ system (Bio-Rad).

### In Vivo Studies

#### Animals and their synchronization

Male and female C57BL/6J mice, 8-12 weeks of age, weighing 18-24 g were used in the in vivo studies with M47. Mice were obtained from Koç University Animal Research Facility (KUTTAM) and experiments were conducted in accordance with the guidelines approved for animal experimental procedures by the Koc University Animal Research Local Ethics Committee (No: 2015/13). Mice were housed in polystyrene cages up to four animals in the room equipped with temperature control (21 ± 2°C) and humidity (55 ± 5%). Mice were housed under 12 h of light (L) in alternation with 12 h of darkness (D) (LD 12:12) prior to any intervention and the same lighting regimen continued to the end of the experiment. Water and food were provided ad libitum throughout the experiments. M47 or vehicle were administered intraperitoneally to mice at the same time every day (three hafter light onset).

#### Preparation of the simple formulation of M47

M47 was dissolved in 2.5% DMSO and 15% Cremophor EL and then diluted with 82.5% isotonic sodium chloride solution (Vehicle = DMSO:Cremophor EL:0.9% NaCl; 2.5:15:82.5, v/v/v) on each study day to prepare freshly, prior to intraperitoneal injection. Solvents were reagent grade, and all other commercially available reagents were used as received unless otherwise stated.

#### Determination of the Dose Range

The dose levels to be used in the single dose toxicity study were selected according to the OECD Guidelines (Guidelines for the testing of chemicals, 2002). Male (n=2-3) and female (n=2-3) C57BL/6J mice were used at each dose level. Mice were treated with 5, 50, 300 or 1000 mg/kg single doses of M47 intraperitoneally, one dose being used per group. Control mice (n=3 for both sexes) were only treated with vehicle (DMSO:Cremophor EL:0.9% NaCl; 2.5:15:82.5, v/v, i.p.). Careful observations of mice including body weight changes, body temperature, behavioral and clinical abnormality, and mortality were performed during 14 days, and gross necropsy of all animals was carried out at the end of the experiment. Body weight was measured every day as an index of general toxicity. M47-induced body weight change was expressed relative to body weight on the initial treatment day. The body temperatures of the mice were recorded by rectal homeothermic monitor (Harvard Apparatus, USA) for 5 days following M47 injection. Temperature measurements were performed at the same time each day. Food intake and water consumption were also monitored for 14 days. After the observation period, mice were exposed to isoflurane anesthesia and blood was collected by cardiac puncture. Mice were immediately sacrificed by cervical dislocation after blood collection. Hematological parameters were analyzed in blood samples.

#### Repeated Dose Toxicity Study-1: Determination of the Maximum Tolerated Dose of M47 Upon 5-Day Administration

The single dose toxicity data were used to assist the selection of the doses in this repeated dose toxicity study. C57BL/6J mice (n=6 per group) were treated with 40, 80 or 150 mg/kg doses of M47 intraperitoneally for 5 days. Control mice (n=4) were only treated with vehicle (DMSO:Cremophor EL:0.9% NaCl; 2.5:15:82.5, v/v, i.p.). Throughout the study, animals were monitored for mortality, clinical signs, body weight changes, body temperature, food and water consumptions, behavior assessment and gross findings at the terminal necropsy. Body weights and body temperatures of mice were measured every day as an index of toxicity. Within 48h after the last dose of M47 administration, blood was collected from mice by cardiac puncture under isoflurane anesthesia. Hematological and biochemical analyzes were performed in blood. Liver, spleen, kidneys and lungs were removed and fixed in 10% formalin solution for histological examinations.

#### Repeated Dose Toxicity Study-2: Determining the Subacute Toxicity of 60 mg/kg M47 Upon 14-Day Administration

According to the results of the 5-day maximum tolerated dose (MTD) determination study, M47 dose was determined as 60 mg/kg for the next 14-day subacute toxicity study. In this study, C57BL/6J mice (n=10) were treated with 60 mg/kg dose of M47 intraperitoneally for 14 days. Control mice (n=5) were only treated with vehicle. Following this, aforementioned procedure (5-day MTD determination study) has been conducted in the same way.

#### Pharmacokinetic Studies

C57BL/6J female mice (n=4 per each time point) were treated with 100 mg/kg i.p. single dose of M47. Blood samples were collected 0, 0.5, 1, 2, 4, 8, 12 and 24h after administration of M47 by cardiac puncture under isoflurane anesthesia. Plasma was obtained by centrifugation from heparinized tubes and stored at −80 °C until analysis. Brain tissues were quickly removed and stored at −80 °C for further processing. For assessment of the brain exposure of M47, levels in this tissue were only determined at 2nd (n=2) and 4th (n=2) haccording to maximum concentrations of M47 was quantified in plasma.

#### Determination of M47 Levels in Plasma and Brain

The Liquid chromatography coupled to tandem mass spectrometry (LC-MS/MS) was used to obtain high mass accuracy data of the analytes in plasma samples for the determination of M47 and internal standard (IS) loratadine. The instrument was operated in full scan mode and ion source parameters were optimized for the analytes. The optimized conditions of the electrospray ionization (ESI) source were as follows: gas temperature, 300°C; gas flow, 10 L/min; nebulizer pressure, 25 psi; capillary voltage, 5500 V. ESI in the positive ion mode provides the best results for M47. For MS-MS detection, the protonated molecules were selected as precursor ions and the most abundant fragment ions obtained at collision energy of 22-20 eV were monitored as the product ions for M47 and IS. MS-MS analysis was performed in the selected reaction monitoring (SRM) positive ionization mode, using mass transitions for M47 *m/z* 472.1 → 353.4 [collision energy (CE) = 22 eV] and Fragmentor 234 V, for IS; *m/z* 383.2→ 337.2 (CE = 20 eV) and fragmentor 140 V. For the separation of the analytes, a conventional reversed-phase LC separation on C_18_-bonded silica was used, with Methanol: % 0.1 Formic acid (90:10, v/v) mobile phase. M47 was extracted from plasma samples by using liquid-liquid extraction with Methanol. 10 µL of sample was injected into the LC-MS/MS system.

#### Pharmacokinetic Analysis

Peak plasma concentration (C_max_) and time to reach peak plasma concentration (t_max_) values were directly obtained from the M47 plasma concentration-time curve. The area under the plasma concentration-time curve from 0 to 48h (AUC_0-48h_) was calculated by the trapezoidal method. AUC_0-∞_ was obtained by addition of the extrapolated part after the last sampling time using standard techniques. Other pharmacokinetic parameters of M47 were calculated by the non-compartmental method. Elimination rate constant (k_el_) was calculated from the terminal points of M47 plasma concentration-time plot and the slope of this line was equal to k_el_. Terminal elimination half-life (t_1/2_) was calculated by ln-linear approximation of terminal points of the data. t_1/2_ and k_el_ were interconverted with the following formula: t_1/2_ = ln2 / k_el_.

#### Survival Study in p53^−/−^ Mice

*p53*^+/-^ mice in C57BL/6J background were purchased from Jackson Laboratory. *p53*^+/-^ mice were bred to obtain *p53*^−/−^ mice. Genotyping of animals were performed provided by Jaxson Laboratory. The experiments with *p53*^−/−^ mice were performed in accordance with the guidelines approved by Koc University Animal Research Local Ethics Committee (No: 2015/13). Animals were maintained in temperature and humidity-controlled rooms under 12 h of light/dark cycle. M47 or vehicle were administered at the same time every day (17 hafter light onset, i.e. HALO17). Food and water were provided *ad libitum*.

Mice were administered 50 mg/kg/day M47 intraperitoneally for 5 days per week, for 7 consecutive weeks (n=9). Control animals were treated with vehicle (n=7). M47 dose was selected according to the results of the repeated dose toxicity studies.

Mortality was recorded daily and Kaplan–Meier survival analysis with log-rank test was performed using GraphPad Prism version 8.00.

#### Statistical Analysis

All data were expressed as means ± standard error of the means (SEM) for each studied variable. Statistical analyses were performed using GraphPad Prism for Windows (GraphPad Software, California, USA). The statistical significance of differences between groups was validated with either of these: Student’s t-test and one- or two-way analysis of variance (ANOVA), following Tukey or Bonferroni post hoc tests, respectively. Type of significant test used in each experiment and their p-values were indicated in the figure legends.

## General

We thank Hiroki Ueda for the generous gift of the *Cry1*^−/−^/*Cry2*^−/−^ mouse embryonic fibroblasts and the *Cry1* rescue vector.

## Author contributions

SG designed the study and performed the experiments and wrote the manuscript. YKA, NaO, and AO perform all mice related experiments. YKA and AO wrote the manuscript. TK, FA and NuO performed apoptosis and gene-editing. SG, SI, SS and ZMG performed in vitro studies. ID and DOU performed pharmacokinetic experiments. ACT handled all mice. OSI and MG analyzed molecule, MT helped computational works. IHK conceived experimental design and wrote the manuscript.

## Data and materials availability

All data needed to evaluate the conclusions in the paper are present in the paper and/or the Supplementary Materials. Additional data related to this paper may be requested from the authors.

## Supplementary Information

**Fig. S1.**
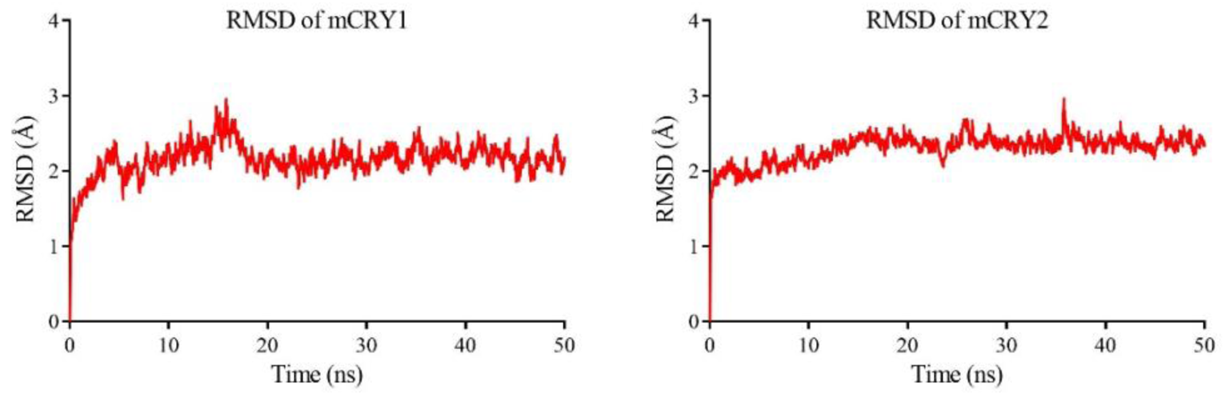
The root-mean square deviation (RMSD) values of CRY simulations. RMSD values of backbone atoms (N-Cα-C) of each amino acid residues were calculated.

**Fig. S2.**
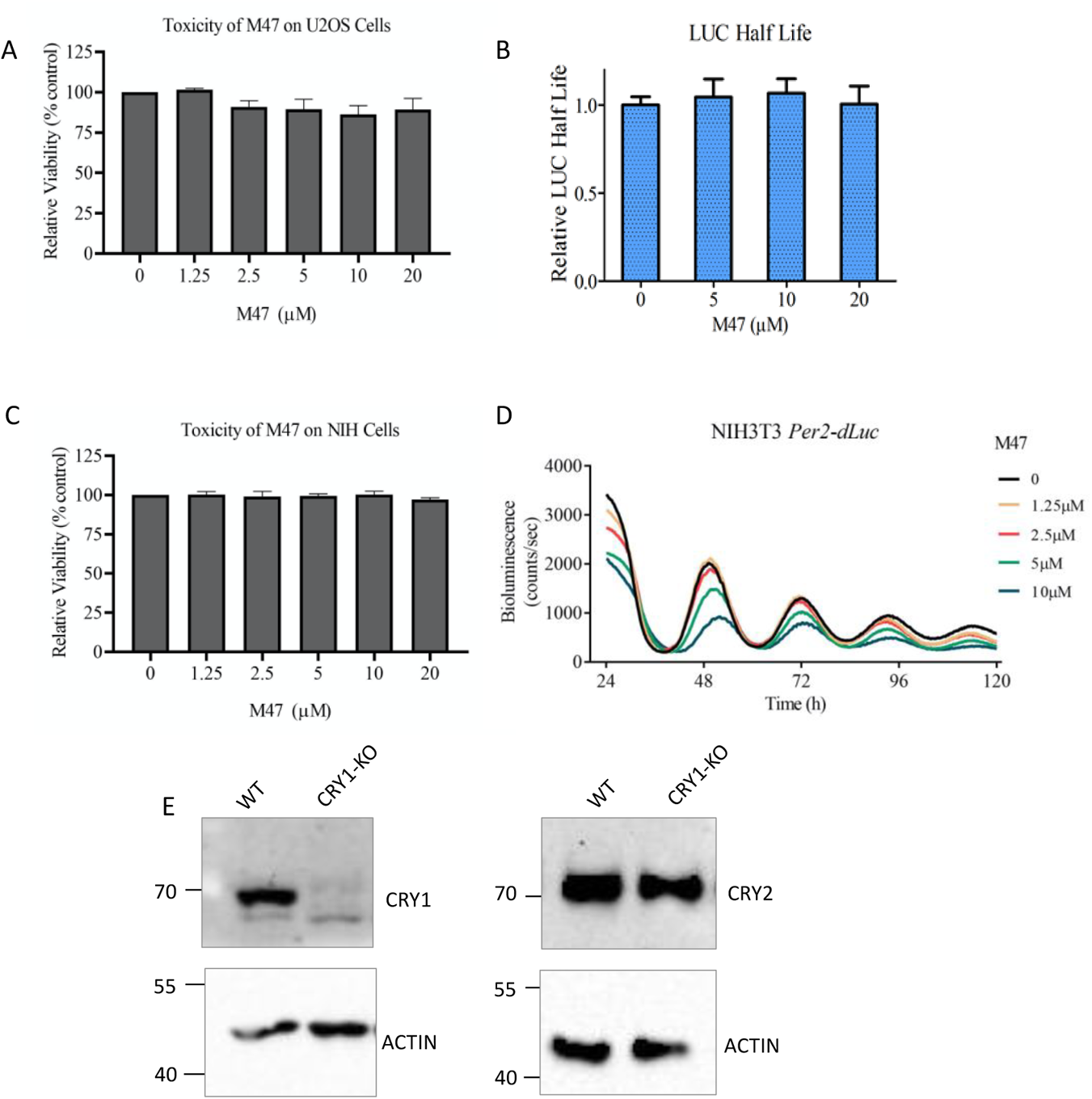
Dose dependent effect of M47 cell viability, LUC degradation and NIH3T3 cells, CRISPR/Cas9 CRY1 deletion confirmation. (A) Cytotoxicity of M47 doses on U2OS cells were tested by the MTT assay (Data represent the mean ± SEM, n=3 with triplicates). (B) M47 did not interfere with the degradation rate of LUC itself. LUC degradation assay was mainly performed as explained in Fig. S3A, however, instead of 40ng *Cry-Luc*, 5ng *Luc* plasmid was transfected due to its high level of expression and luminescence (Data represent the mean ± SEM, n=3 with triplicates). (C) Cytotoxicity of M47 doses on NIH 3T3 cells were tested by the MTT assay (Data represent the mean ± SEM, n=3 with triplicates). (D) Dose dependent effect of M47 was tested on the NIH3T3 cells stably expressing *Per2-dLuc* (Representative luminescence data of three independent experiments (n=3) with duplicates). (E) Confirmation of CRY1 knockout in the U2OS cell line. Immunoblot of CRY1 and CRY2 proteins in the U2OS clone was used to confirm that specific knockout of CRY1 was successful. Actin was blotted as the loading control. Numbers on the left of each panel indicate the positions of the corresponding molecular size markers in kDa. WT: Wildtype; KO: CRY1 knockout.

**Fig. S3.**
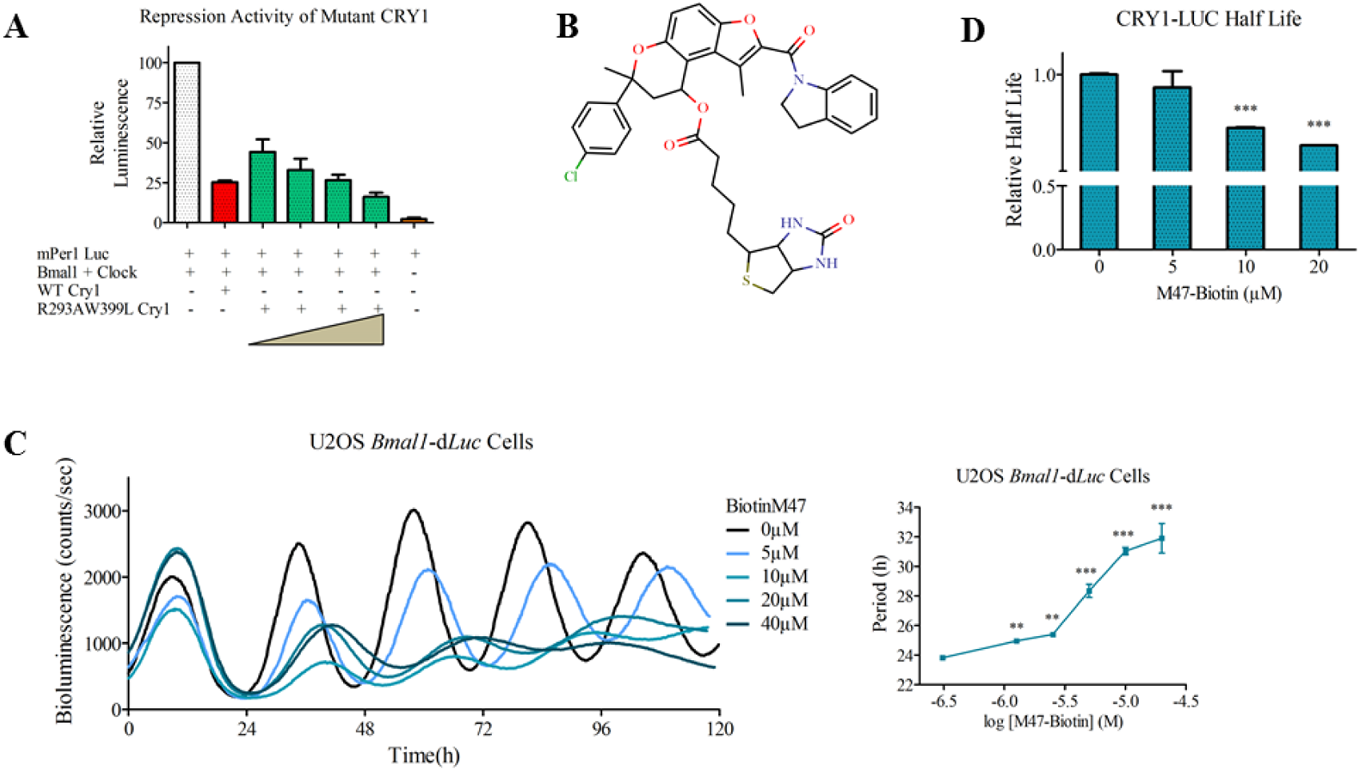
Activity of mutant CRY and effect of biotin on the activity of M47. (**A**) Mutant R293AW399L CRY1 was active and could repress the BMAL1:CLOCK mediated transcription. Wildtype CRY1 was used as the positive control. Data were normalized according to transcription activity of BMAL1:CLOCK without ectopic expression of CRY1 (Data represent the mean ± SEM, n=3 with triplicates). (**B**) Chemical structure of bM47 used for pull-down assays. (**C**) Biotin attached to M47 did not change the effect of M47 on the circadian rhythm. Luminescence data is a representative of two independent (n=2) experiments with duplicates. Period data is reported as the mean ± SEM, n=2 with duplicates ***: p < 0.001 **: p < 0.005, * : p < 0.05 versus DMSO control by student t-test. (**D**) bM47 enhanced the degradation rate of CRY1-LUC. Data represent the mean ±SEM, n=3 with triplicates ***: p < 0.001 versus DMSO control by student t-test.

**Fig. S4.**
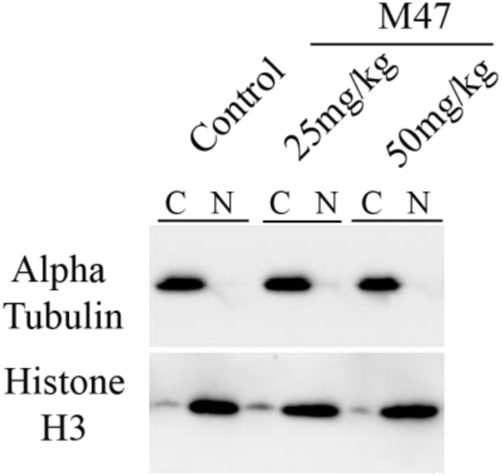
Subcellular fractionation control of mice liver. Liver samples were fractionated as cytosolic (C) and nuclear (N) parts. Alpha tubulin and Histone H3 were probed as the cytosolic and nuclear protein markers, respectively.

**Table S1.**
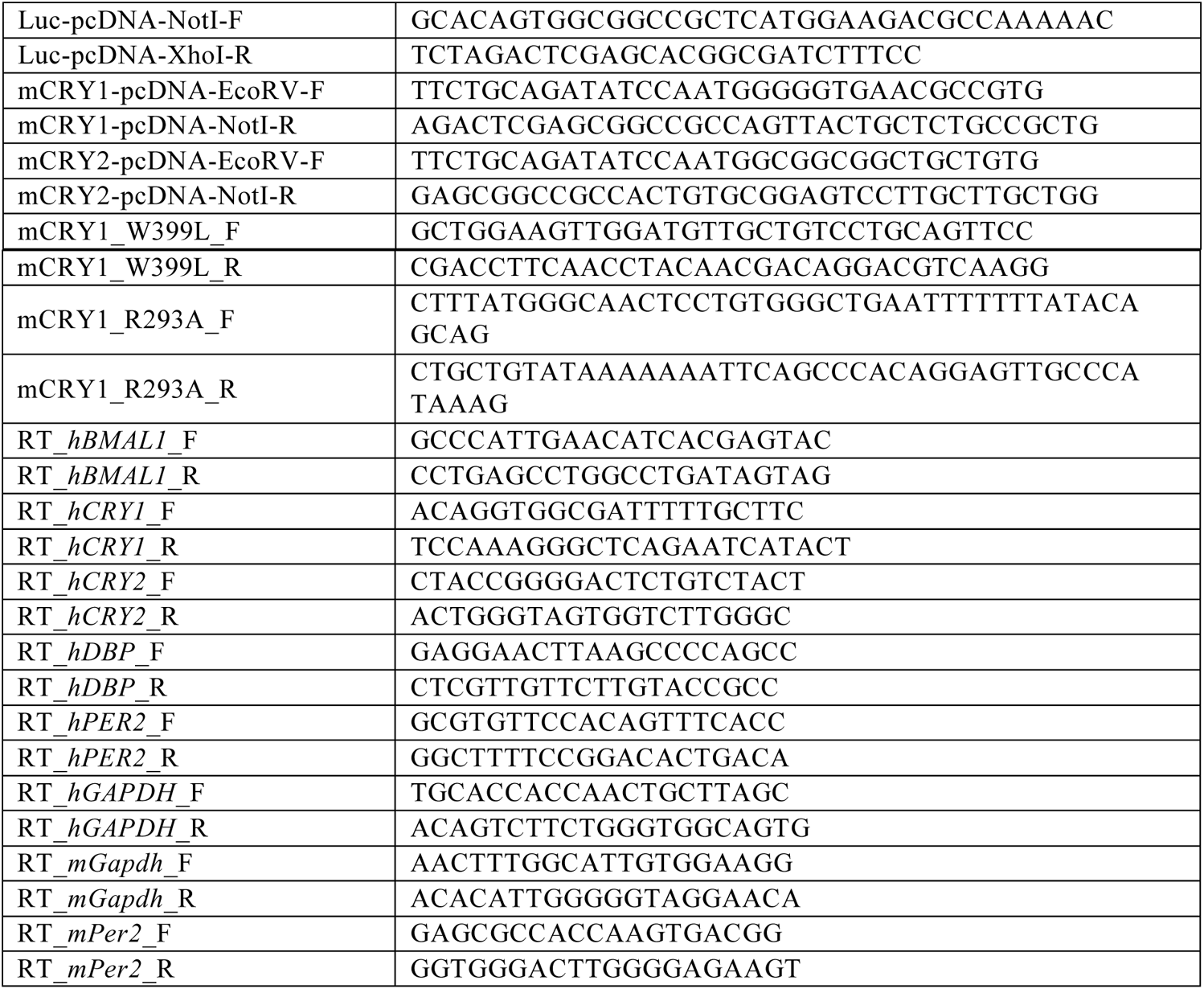
Sequence of primers for Cry-Luc cloning, site-directed mutagenesis and RT-qPCR.

